# A Two-Dimensional Grid-Cell Code for Three-Dimensional Navigation in Freely Flying Bats

**DOI:** 10.64898/2026.06.10.728358

**Authors:** Kevin K. Qi, Michael M. Yartsev

## Abstract

Animals evolved to navigate three-dimensional worlds, a capacity that relies on internal representations of position. In one- and two-dimensional environments, grid cells in the medial entorhinal cortex provide a periodic signal for self-location that persists throughout the environment and is thought to arise from toroidal population dynamics. Yet global grid periodicity has not been observed during navigation through three-dimensional volumes, leaving unresolved whether grid cells and the underlying two-dimensional toroidal code can support navigation in three dimensions. Using large-scale wireless recordings in freely flying bats, we found that grid cells retained periodic firing during flight and that co-modular ensembles exhibited toroidal organization across behavioral states. Analyses of flight geometry in the laboratory and the wild showed that bats largely traverse volumetric space along two-dimensional planes of motion. Toroidal phase and grid-cell firing patterns mapped onto these planes, revealing how a two-dimensional neural code aligns with behaviorally relevant subspaces to support navigation in a three-dimensional world.

## Introduction

Spatial navigation relies on internal representations of position^1,2^. In mammals, a key component of this system are grid cells in the medial entorhinal cortex (MEC), whose firing fields form a periodic lattice in two-dimensional (2D) environments^1,3–5^. In rodents moving in one- or two-dimensional environments, the spatial periodicity of grid cells is thought to arise from population dynamics evolving on a low-dimensional toroidal manifold, as formalized in continuous attractor network (CAN) models^1,6–17^. Yet, whether and how such computations extend to three-dimensional (3D) environments remains unresolved. Although many animals live and navigate in 3D space, globally periodic grid cells have not been observed during navigation through 3D volumes^18,19^, and it remains unknown whether the toroidal network dynamics described in rodents generalize to other species or to 3D navigation. Even if toroidal grid-cell organization is conserved across species, it remains unclear how such a two-dimensional code can support navigation in a three-dimensional environment. A toroidal code provides a natural periodic coordinate system for position on a 2D plane^20–22^, but a 3D volume contains an infinite number of possible planes. A toroidal code thus faces an apparent dimensionality mismatch: a two-dimensional code cannot inherently represent position in three dimensions without an additional, dimensionality-reducing principle that is presently unknown.

Here we addressed these gaps by wirelessly recording large-scale ensemble activity from the medial entorhinal cortex of freely flying bats (*Rousettus aegyptiacus*) during spontaneous, unconstrained three-dimensional spatial foraging behavior. This approach allowed us to directly test (i) whether periodic firing of grid cells is preserved during three-dimensional movement, (ii) whether the population dynamics are consistent with an underlying toroidal organization in a species specialized for navigation in three dimensions, and (iii) how a two-dimensional neural code could be mapped onto movement through a three-dimensional world.

## Results

### Periodic grid cells in flying bats

To study grid cells from the single-cell to the ensemble level during unconstrained 3D navigation, we used Neuropixels to wirelessly record the simultaneous activity of hundreds of single units from the MEC of freely flying bats engaged in spontaneous aerial foraging^23–25^ (*n* = 3 bats, 13 sessions; Fig. 1A-D). During each recording session (mean duration: 1 hour 55 min), bats flew a diverse repertoire of flight paths (74 ± 23 flights, mean ± s.d. per session) throughout the 3D volume between automated feeder platforms and self-selected perch locations (Fig. 1A and Fig. S1). Many of these flight paths were repeated with high precision^24^, and were thus termed ‘clustered flights’ (Fig. 1B; 662 of 968 flights, 68.4%, across all bats). Flight paths not repeated within the span of a recording session were termed ‘unclustered flights’ (306 of 968 flights, 31.6%). During these foraging sessions, we recorded a total of 2,353 well-isolated single units (181 ± 111, mean ± s.d. per session) from the superficial layers II/III of MEC. Consistent with previous reports^18,19^, when we examined the activity of the single units as a function of self-position by pooling together all flights, we did not observe global 3D lattice structure beyond chance levels (Fig. S2). Yet, when we examined the same neural activity along the bats’ self-selected flight trajectories, robust periodic firing emerged (Fig. 1E,F). We identified 1,102 units that exhibited multiple, stable firing fields on clustered flight paths (Fig. S3) and asked whether MEC activity was organized not only by *where* the animal was (its 3D position), but also by *how* it chose to traverse the 3D environment (the specific path taken). Indeed, periodic structure was largely obscured when single-unit activity was aligned only to take-off, without accounting for the bat’s trajectory (Fig. S4A). When the same flights were instead grouped by trajectory cluster, periodic activity was evident along the full extent of each repeated flight path (Fig. 1E,F, and Fig. S4B). The pattern was consistent within each flight cluster but differed between clusters.

**Figure 1.**
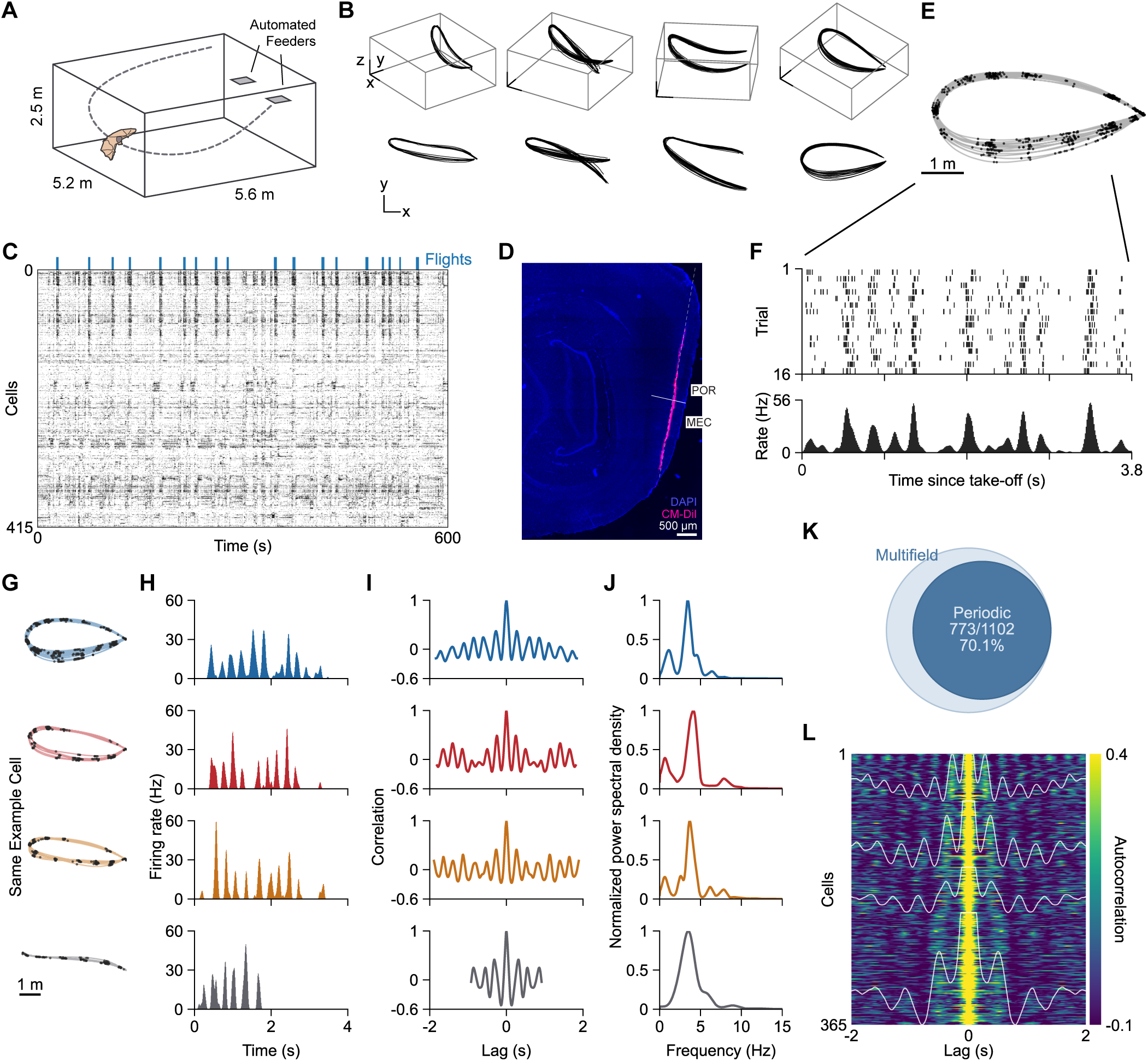
Periodic grid cells in flying bats. **(A)** Schematic of flight enclosure with illustrative bat flight trajectory (dashed line) and automated feeder platforms (gray squares). **(B)** Four example clustered flight paths (columns) shown in 3D perspective (top), and top-down projection (bottom). Scale bar 1 m. **(C)** Rastermap^74^ visualization of the neural activity of 415 simultaneously recorded cells during a 600-s segment of a recording session. Blue bars indicate flights. **(D)** Sagittal section from one recorded bat showing Neuropixels 1.0 probe track through the medial entorhinal cortex (MEC; white dashed line), stained for 4′,6-diamidino-2-phenylindole (DAPI, blue) and CM-DiI (pink). POR, postrhinal cortex. Data were collected from the probe contacts located in MEC. **(E)** Spikes from an example MEC neuron (black) overlaid on flight trajectories (gray). **(F)** Top, raster plot of the neuron in **E** aligned to flight take-off. Bottom, trial-averaged firing rate aligned to take-off. **(G)** Spikes of an example grid cell (black dots) across four different flight trajectories (colored). Note the conserved periodic firing across the different flight trajectories. **(H)** Trial-averaged firing rate of the grid cell in **G** aligned to take-off. **(I)** Autocorrelation of the trial-averaged firing rate in **H**. **(J)** Peak-normalized power spectral density of the firing rate modulation shown in **H**, revealing conserved periodic structure. **(K)** Percentage of multifield cells classified as significantly periodic. **(L)** Time-lagged firing rate autocorrelograms of significantly periodic grid cells, clustered across sessions by their power spectral profile and sorted by mean cluster grid spacing. White traces indicate mean autocorrelation of each cluster. Only clusters with 50 cells or more are shown.

Because all flights were aligned to take-off, these cluster-specific patterns indicate that periodicity tracked the animal’s position along its path rather than elapsed time. Notably, while field locations differed between paths, the periodicity of each unit was conserved across them (Fig. 1G). Leveraging the natural trial structure that emerged spontaneously during foraging, in which different clustered flights were repeated with high precision, we quantified the firing periodicity using the autocorrelation and power spectral densities (PSD) of trial-averaged linearized firing rates (Fig. 1H-J). More than 70% of the multifield units exhibited robust periodic firing along clustered flight paths (773/1,102 or 70.1%; Fig. 1K). Moreover, clustering of PSD profiles revealed groupings of units with similar grid spacing (Fig. 1L), arranged with increasing scale along the dorsal-to-ventral axis of MEC (Fig. S5), reminiscent of the modular organization of grid cells observed in rodents moving in 2D environments^26^. Together, these findings show that global grid-cell periodicity is preserved during unconstrained, spontaneous movement in three-dimensional space but is revealed only when neural activity is evaluated with respect to the flight paths that bats naturally select.

### Toroidal topology of grid-cell population activity in flying bats

In rodents, co-modular grid-cell activity exhibits toroidal topology in 1D and 2D environments consistent with CAN models^1,6–9,13,14,17^. Yet it remains unknown whether such toroidal organization generalizes across species, and whether it persists during three-dimensional movement. We therefore examined the population activity of co-modular grid cells in flying bats. First, we divided each session into flight bouts, including both clustered and unclustered flights, and non-flight bouts. Then, leveraging the stable nature of co-modular grid-cell spike-time relations^10–17,27,28^, we identified grid modules using spike-time-correlation-based clustering^13,14,17^, yielding five modules that exhibited stable within-module spike-time correlations across both flight and non-flight behavioral states (Fig. 2A,B, and Fig. S6; Methods). We used persistent cohomology to identify long-lived topological features in the high-dimensional activity of co-modular grid cells, summarized in a barcode of feature lifetimes^13,14,17,29,30^. Consistent with the toroidal topology reported for population activity in the rodent MEC^13,14,17^, persistent cohomology revealed that the population activity of these modules exhibited the prominent features expected for a two-dimensional torus: a single connected component, two circular dimensions, and one enclosed cavity (Fig. 2C, Fig. S7, and Table S1; Methods). Furthermore, we reasoned that if the spatial activity of grid cells indeed reflects attractor dynamics on the internal toroidal manifold, then units with similar toroidal tuning should also exhibit more similar spatial firing patterns during flight. To test this, we used cohomological decoding^13,14,17,29^ to construct a coordinate system on the toroidal manifold, with each circular dimension parameterized by an angular value (Methods). We then computed rate maps of each grid cell with respect to both external Euclidean coordinates (spatial rate maps; Fig. 2D, left) and internal toroidal coordinates (toroidal rate maps; Fig. 2D, middle and right, for flight and non-flight, respectively; Fig. S8A). Pairwise similarity of co-modular grid cells was strongly correlated between their spatial and toroidal rate maps (Fig. 2E,F; *r* = 0.56 ± 0.11, mean ± s.d. across modules; all modules: *P* < 0.001; see also Fig. S8B,C), suggesting that grid cells which exhibit similar spatial firing patterns are also located nearby on the toroidal manifold, as predicted from CAN models and consistent with experimental data in rodents^1,6–17,27^. Finally, if grid cells operate on an intrinsic toroidal manifold, this population structure should persist across behavioral states^1,6–17,27^, including states in which the animal is not moving. We therefore asked whether toroidal tuning was preserved between flight and non-flight periods, during which bats were stationary, including while hanging upside down. The tuning of grid cells with respect to toroidal coordinates remained highly stable across behavioral states (Fig. 2D,G,H; *r* = 0.92 ± 0.04, mean ± s.d. across modules; all modules: *P* < 0.001; see also Fig. S8D; Methods) and toroidal coordinates decoded from the non-flight periods closely matched those decoded directly from flight bouts (Fig. S8E,F; Methods). Together, these findings indicate that the activity of co-modular MEC populations in freely flying bats is consistent with an underlying persistent two-dimensional toroidal organization, similar to that observed in rodents^13,14,17^, despite the animals’ navigation through three-dimensional space.

**Figure 2.**
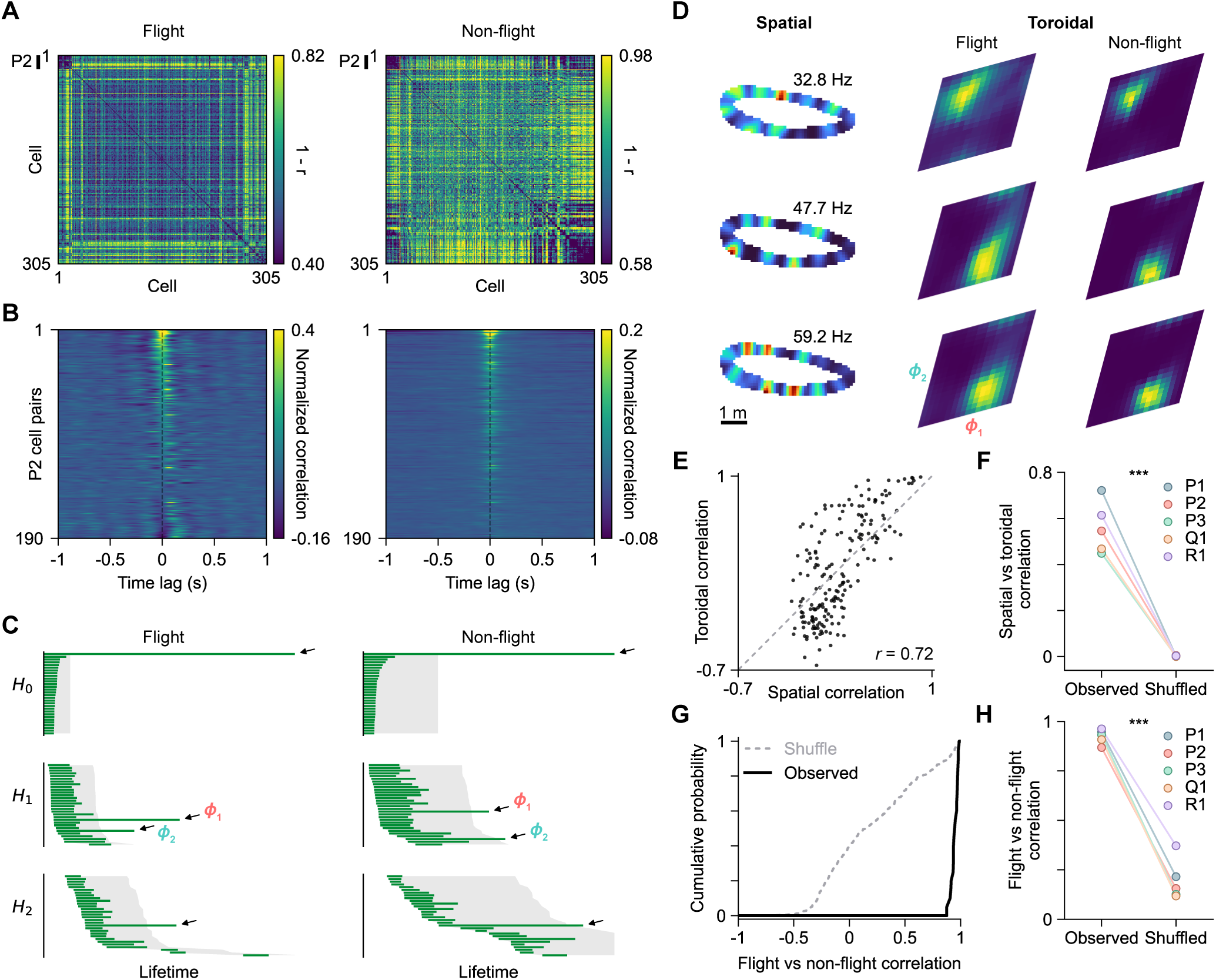
Toroidal topology of grid-cell population activity in flying bats. **(A)** Pairwise correlation distance matrix between simultaneously recorded cells during all flight (left) and all non-flight (right) periods. Both flight and non-flight distance matrices are sorted identically according to grid module clustering. P2 indicates a putative grid module (bat P, module 2). **(B)** Normalized cross-correlograms for all cell-pairs of the module shown in **A** during flight (left) and non-flight (right), ordered by zero-lag correlation measured during flight. Cross-correlations were normalized by mean-subtraction. **(C)** Persistent cohomology analysis of co-modular P1 population activity during flight (left) and non-flight (right). Barcodes (green) show one long-lived connected component (H0), two one-dimensional cycles (H1), and one two-dimensional cavity (H2), consistent with a two-dimensional torus. Gray shaded area indicates 95^th^ percentile significance thresholds. Arrows denote significant topological features of each dimensionality. **(D)** Spatial rate maps and toroidal rate maps (peak-normalized) of three example grid cells from P2. Spatial rate maps are shown on the left. Rate map peak firing rate is indicated. Toroidal rate maps are shown in the middle for flight bouts and right for non-flight bouts. **(E)** Pairwise spatial correlation (between spatial rate maps) versus pairwise toroidal correlation (between toroidal rate maps during flight) of P1 grid-cell pairs (*n* = 190 pairs, Pearson *r* = 0.72). Gray dashed line, unity line. **(F)** Pearson correlation between pairwise spatial correlation and pairwise toroidal correlation for each of the five grid modules compared to shuffle distribution constructed by random permutation of unit identity. *n* = 190 (P1), 190 (P2), 210 (P3), 378 (Q1), 253 (R1) cell pairs. \*\*\**P* < 0.001. **(G)** Observed and shuffled (black and gray, respectively) cumulative distribution of pairwise correlation between toroidal rate maps during flight and non-flight for the P1 module. Shuffled distribution is constructed by random permutation of unit identity. **(H)** Mean within-module pairwise toroidal rate map correlation during flight and non-flight bouts for each of the five different grid modules versus shuffle as constructed in **G**. *n* = 20 (P1), 20 (P2), 21 (P3), 28 (Q1), 23 (R1) cells. \*\*\**P* < 0.001.

### Mapping of grid-cell representations on the plane of motion

The two-dimensional toroidal topology of grid-cell population activity raises a fundamental paradox: how can an intrinsically two-dimensional internal code support navigation through a three-dimensional world? A toroidal manifold provides a natural coordinate system for position on a two-dimensional plane^20–22^, but volumetric space does not define a unique plane onto which this code should be mapped. We therefore asked whether the missing reference geometry might arise not from the environment alone, but from the structure of the animal’s own movement within it. To address this, we first examined the ethology of bat flight and asked how bats naturally move through three-dimensional environments, both in the wild and in the laboratory. In the wild, Egyptian fruit bats perform nightly commutes extending over dozens of kilometers to remote foraging sites^31–33^. We analyzed GPS tracking data from such long-range outdoor flight^34^ and observed that trajectories were organized into a sequence of distinct movement epochs: (i) an ascent towards a cruising altitude, (ii) extended travel at a relatively stable elevation, and finally (iii) descent towards the foraging site (Fig. 3A,B). Each epoch (ascent, cruise, and descent) was well described by a single plane of motion (Fig. 3C). Thus, the long-range commuting flights of wild Egyptian fruit bats can be described as directional movement through three-dimensional space unfolding across successive planes of motion.

**Figure 3.**
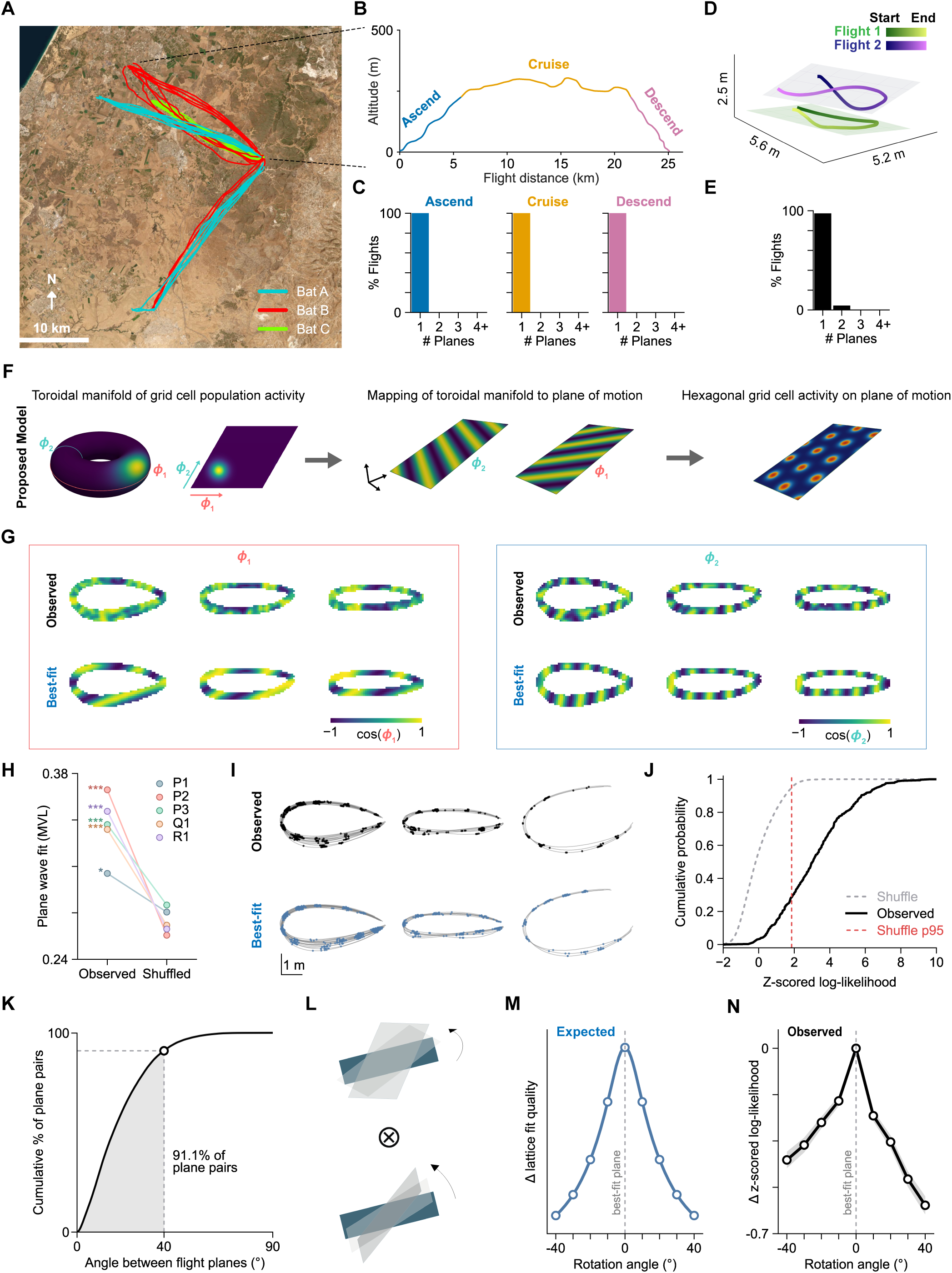
Mapping of grid-cell representations on the plane of motion. **(A)** Flight trajectories of three example Egyptian fruit bats (colors) over multiple nights in the wild (individual traces). **(B)** Altitude profile of a representative commute flight, showing ascent (blue), cruise (orange) and descent (pink) phases. **(C)** Distribution of number of planes required to explain >98% of positional variance during ascent, cruise, and descent phases (*n* = 75 flights from 13 bats). Most segments were captured by a single plane. **(D)** Two example flight paths during spontaneous foraging in the laboratory flight room. Line color indicates progression from start to end. **(E)** Distribution of number of planes required to explain >98% of the positional variance for laboratory flights. Most flights were captured by a single plane (96.5% or 934/968 of flights). **(F)** Proposed plane-referenced grid model. A two-dimensional toroidal manifold (left, shown as folded and unfolded) is mapped onto the plane of motion of each flight, such that toroidal axes (***ϕ***1 and ***ϕ***2) generate plane-wave components (middle) whose combination gives rise to hexagonal structure on that plane (right). **(G)** Top, observed toroidal phase maps of ***ϕ***1 (left, red box) and ***ϕ***2 (right, blue box) of the P2 module on clustered flight paths in the plane of motion. Bottom, best-fit plane-waves on the plane of motion. **(H)** Mean vector length (MVL) quantifying the plane-wave fit for each module, compared to shuffled data. Points denote modules (color-coded); lines connect observed and shuffled values. *n* = 20 (P1), 20 (P2), 21 (P3), 28 (Q1), 23 (R1) cells. \*\*\**P* < 0.001. \**P* = 0.023. **(I)** Top, observed spikes (black) overlaid on flight trajectories (gray) for an example cell. Bottom, corresponding best-fit spike positions (blue) by the plane-referenced grid model on the plane of motion, overlaid on the same flight trajectories. **(J)** Cumulative distribution of z-scored log-likelihood values for hexagonal lattice fits (black) compared with that of shuffle controls across the population (gray). Red dashed line indicates 95^th^ percentile of z-scored empirical shuffle distribution. Note, significance is determined per cell using the empirical P value rather than the *z*-score. The shuffle population 95^th^ percentile is provided as a reference and not used for the determination of per-cell significance. **(K)** Cumulative distribution of inter-plane angles, with 91.1% (shaded region) of plane pairs within 40° of each other. **(L)** Schematic of candidate planes generated by rotating the plane of motion around the azimuth and pitch axes. **(M)** Expected change in lattice-fit quality as candidate planes are rotated away from the plane of motion, defined as the best-fit plane to the bat’s flight path. If grid-cell representations are aligned to the plane of motion, lattice-fit quality should be maximal at 0° rotation and decrease monotonically with increasing angular deviation. **(N)** Observed mean difference between z-scored log-likelihood on rotated flight planes and best-fit plane of motion (average slope of lattice-fit reduction *β* = -0.139 ± 0.008 (mean ± s.e.m.) *z*-score / 10°; Wilcoxon signed-rank, *W* = 2279, *n* = 283 cells, *P* = 3.14 × 10^−38^). Gray shaded region denotes s.e.m. of the mean trace.

These GPS tracking data resolve the kilometer-scale structure of commute flights but lack the spatial and temporal resolution needed to capture the fine-scale geometry of foraging^31,32^. We therefore examined the geometry of flight during spontaneous foraging in the laboratory, where bats express comparable meter-scale foraging behavior^23–25^ and trajectories can be tracked at centimeter and millisecond resolution, and include non-trivial geometries such as highly curved loop flights. Here too, a planar movement pattern persisted. While the three-dimensional flight trajectories of bats varied widely (Fig. 1B) and occurred throughout the flight room (Fig. S1), nearly all individual trajectories, both clustered and unclustered, were well described by a single plane of motion that accounted for the vast majority of spatial variance within the flight path (Fig. 3D,E). These planes spanned a range of orientations (Fig. S9): many were near horizontal, but a substantial subset was tilted away from it, and their relative orientations varied across trajectories. Thus, although bats moved through a three-dimensional volume, their flight repertoire was not confined to a single two-dimensional subspace, but decomposed into multiple trajectory-specific planes of motion with different orientations, a natural behavioral substrate onto which grid-cell representations could be mapped.

We therefore hypothesized that the dimensionality mismatch between a two-dimensional grid code and three-dimensional space could be resolved by mapping grid-cell representations onto the animal’s present plane of motion (the two-dimensional subspace defined by the top two principal components of each flight trajectory), a framework we term a plane-referenced grid code (Fig. 3F). Unlike earlier work on planar encoding on locomotor surfaces^35–37^, where the relevant plane was defined by the surface itself, here the plane is self-selected and varies across trajectories. Importantly, this framework makes direct predictions at both the population and single-cell levels. At the population level, the toroidal manifold, inferred from population activity rather than flight geometry and preserved during non-flight periods when bats were stationary, should align with the self-selected plane of motion. Under this alignment, the two circular coordinates of the torus would correspond to plane-wave components of a hexagonal lattice on that plane^7,8,13–15^ (Fig. 3F, middle, and Fig. S10A-C). At the single-cell level, firing along individual flight trajectories should be consistent with an underlying hexagonal lattice defined on the corresponding trajectory-specific flight plane^15,27^ (Fig. 3F, right, and Fig. S10D).

To test these predictions, we first estimated toroidal phase maps by computing the mean toroidal phase (***ϕ****1* and ***ϕ****2*) at each position during flight (Fig. 3G; Methods). If the toroidal manifold coordinates are aligned with the flight plane, phase patterns along each flight path should match those resulting from the intersection of the trajectory with plane waves defined on the plane of motion. Consistent with this prediction, plane-wave fits to the toroidal phase maps of each module revealed that the observed phase patterns were well captured by plane waves defined on the flight plane (Fig. 3G,H; mean vector length (MVL) = 0.34 ± 0.02, mean ± s.d. across modules; P2, P3, Q1, R1: *P* < 0.001; P1: *P* = 0.023; Methods). We next asked whether the firing patterns of individual grid cells were consistent with a hexagonal lattice on the plane of motion. To do so, we computed the log-likelihood of observed spike positions under candidate hexagonal grid patterns defined on the plane of motion, spanning the full range of grid spacings, orientations, and phases. Because this approach operates directly on raw spike positions, it avoids the need for additional hyperparameters for rate map estimation but is sensitive to background and out-of-field spiking activity. We therefore restricted the analysis to cells with a minimum of two prominent peaks in their inter-spike distance distribution (*n* = 331 cells). Over 70% of these cells exhibited firing patterns consistent with a hexagonal lattice on the plane of motion (*n* = 236/331, 71.3%; example cell, Fig. 3I and Fig. S11; population, Fig. 3J; Methods).

To further assess the specificity of this mapping onto the animal’s actual plane of motion, relative to other plane orientations the bats ethologically occupy (Fig. S9), we repeated the grid-fitting analysis on 48 nearby candidate planes generated by rotating the plane of motion up to 40°, spanning the range of relative plane orientations the bats naturally adopted (Fig. 3K), in 12 azimuthal directions (Fig. 3L). Each trajectory’s plane of motion was defined from the flight path alone, independently of neural activity. If grid-cell representations are aligned to the plane the animal is currently traversing, lattice-fit quality should be highest on the observed plane of motion and deteriorate monotonically as candidate planes are rotated away from it (Fig. 3M). Consistent with this prediction, the lattice-fit quality of grid cells was maximal on the true plane of motion and declined monotonically with increasing rotation angle (Fig. 3N, and Fig. S12A-C). Spatially tuned multifield non-grid cells did not exhibit a reduction in lattice-fit quality with increasing rotation angle (Fig. S12D), as further confirmed with simulations of grid cells and non-grid multifield cells (Fig. S12E), indicating that this specificity was not a general property of spatially tuned cells. Together, these results suggest that grid-cell periodicity and toroidal tuning during clustered flights are specifically aligned to the animal’s present plane of motion.

The specificity of this mapping raised a further prediction across flight planes: if an internally organized toroidal grid code maps onto each flight plane, then aligning activity from different flight planes in a shared grid coordinate system should recover a common hexagonal lattice. This imposes an additional constraint relative to the single-plane fits above, since one lattice must account for activity across multiple flight planes rather than on each plane separately. Because each flight sparsely samples its trajectory-specific plane, limiting recovery of a two-dimensional lattice, we restricted this analysis to periodic cells with greater spatial and spike sampling (≥300 spikes on clustered flights from recording sessions with at least three long [≥8 m] flight clusters; n = 136 cells). In room-referenced coordinates, gridness scores, which quantify the six-fold rotational symmetry of rate-map autocorrelations (Fig. 4A and Fig. S13; Methods), did not exceed the shuffle expectation, and the fraction of cells with significant gridness (9 of 136, 6.6%) did not exceed the 5% expected by chance (binomial P = 0.24). However, after alignment in shared plane-referenced coordinates, gridness was significantly higher than in room-referenced coordinates, beyond the increase expected from aligning shuffled data (per-cell difference in z-scored gridness, Wilcoxon signed-rank test, W = 1310, P = 3.53 × 10^−13^; Fig. 4A,B), and significant gridness was detected in 46 of 136 cells (33.8%; binomial P = 6.99 × 10^−26^; McNemar exact P = 3.02 × 10^−9^ versus room-referenced coordinates). Thus, aligning activity across trajectory-specific flight planes revealed a shared hexagonal grid representation that was obscured in room-referenced coordinates.

**Figure 4.**
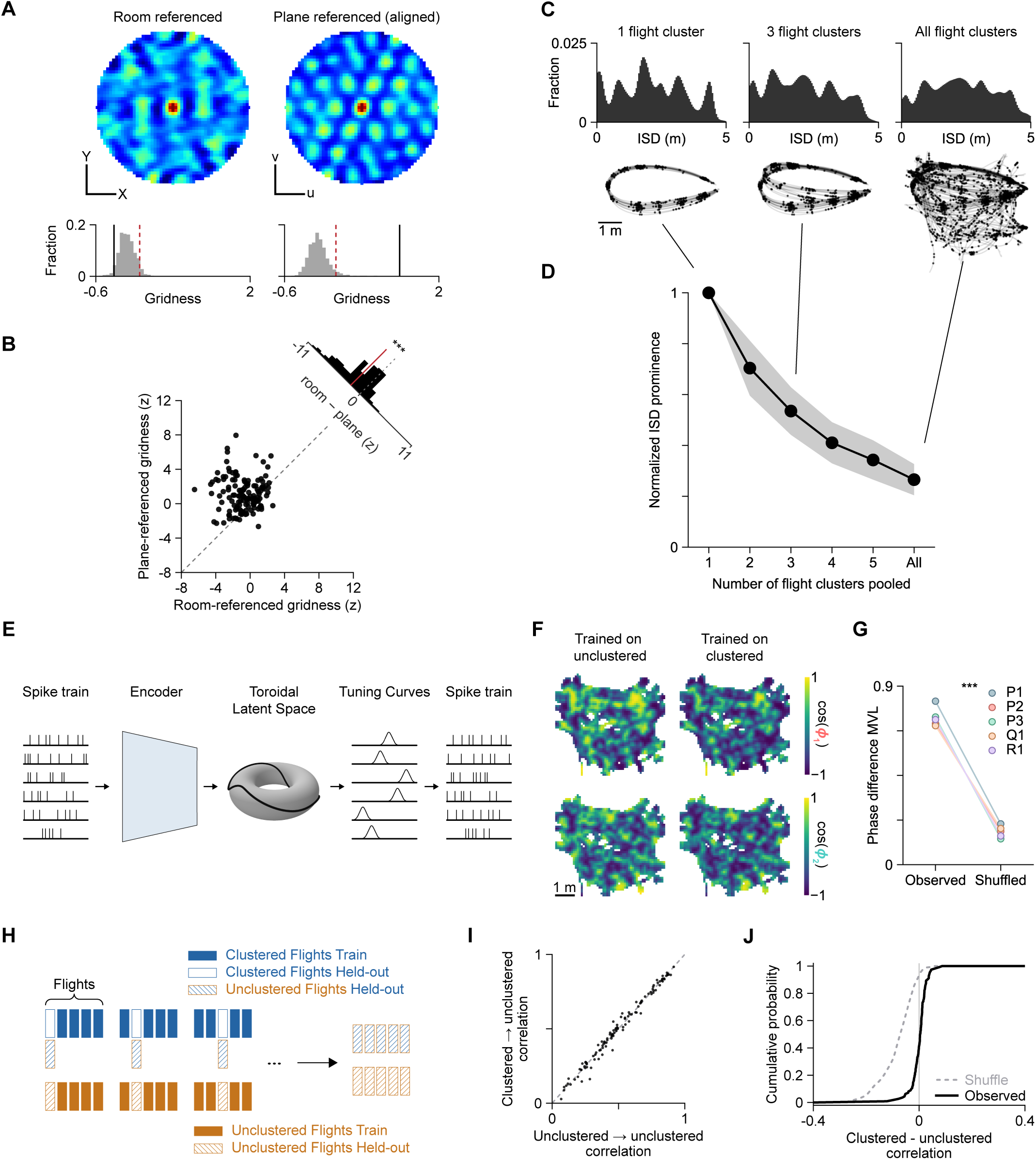
Grid-cell representations generalize across flight planes and to unclustered flights. **(A)** Spatial rate-map autocorrelations of an example cell computed in the room-referenced top-down projection (XY-plane, left column) and shared plane-referenced coordinate system (uv-plane, right column). Color scale, -0.3 to 0.85. Scale bar, 1 m. Bottom, distribution of shuffled gridness scores for the same cell with the corresponding observed gridness score (black) relative to the 95^th^ percentile significance threshold (dashed red). **(B)** Gridness of each cell (*n* = 136) in room-referenced versus plane-referenced coordinates, expressed as a z-score relative to the cell’s shuffle distribution in each frame. Dashed line, unity. Top right, distribution of per-cell differences (room-referenced minus plane-referenced); red line, mean difference. Wilcoxon signed-rank test, W = 1310, ***P = 3.53 × 10^−13^. **(C)** Example recorded grid-cell response obtained by pooling one (left), three (middle), and all (right) flight clusters. Top, inter-spike-distances (ISD) of between-flight spike pairs. Bottom, spikes (black) on flight trajectories (gray). **(D)** Lattice structure, as measured by the peaks in the ISD using a normalized prominence score (black trace, median; Methods) as increasing numbers of flight clusters are combined. Shaded region indicates the bootstrapped standard error of the median (n = 197 cells). **(E)** Schematic of the latent variable model (LVM) with toroidal latent space, based on the architecture proposed by Bjerke et al.^38^. **(F)** Example top-down projected toroidal phase maps of the P2 module showing ***ϕ***1 (top) and ***ϕ***2 (bottom) during unclustered flights decoded from LVMs trained on unclustered flights (left) and on clustered flights only (right). Scale bar 1 m. **(G)** Mean vector length (MVL) of the angular difference between toroidal phase maps decoded by LVMs trained on unclustered versus on clustered flights for the five grid modules, taken as the median across 100 training seeds. Shuffled values are computed by circularly shifting the decoded toroidal coordinates within each unclustered flight trial by a random offset. \*\*\**P* < 0.001. **(H)** Illustration of the 5-fold train-test procedure used to compare held-out single-cell activity of LVMs trained only on clustered flights (blue) with LVMs trained only on unclustered flights (orange). For each fold, LVM was trained on 80% of either the clustered flight data (blue solid bars) or unclustered flight data (orange solid bars), then both were evaluated on 20% of the unclustered flight data (striped bars). **(I)** Paired scatter plot of per-cell (*n* = 112) Pearson correlation between predicted and observed firing rates on held-out unclustered flights using LVM trained on clustered flights only (y-axis) and on unclustered flights only (x-axis). Clustered minus unclustered correlation difference, 0.0013 ± 0.030 (mean ± s.d.). Dashed line indicates unity line. **(J)** Cumulative distribution (black) of the per-cell Pearson correlation differences of the paired comparisons in **I**. Dashed curve indicates correlation difference using shuffled data, where a random circular shift was applied to each cell’s spike train during clustered flights (Mean observed correlation difference minus shuffled correlation difference = 0.083, *n* = 100 shuffles, *P* < 0.01).

While the plane-referenced grid framework accounts for the observed periodicity within and across clustered flights, we further asked whether it could also explain neural activity on unclustered flights, where periodicity was not apparent (Fig. S4). The framework makes three predictions. First, because periodicity is expressed relative to each flight’s own plane, pooling flights occurring on different planes should progressively degrade apparent periodicity. Consistent with this prediction, grid periodicity decreased monotonically as flight paths on distinct planes were progressively pooled (Fig. 4C,D), and this reduction was reproduced by simulations of plane-referenced grid cells in which each individual flight was periodic by construction (Fig. S14A,B). Second, if grid cells are referenced to the movement plane, their room-referenced spatial stability should be reduced on unclustered flights. While stability of grid cells was indeed lower on unclustered flights, they remained above that expected from random alignment of grids onto flight planes, suggesting the possibility of non-random plane-referencing (Fig. S14C-F). In line with this notion, we found that the activity on the shared flight segments between different flight clusters was highly similar (normalized stability = 0.93 ± 0.03, median ± s.e.) when traversed in the same direction (Fig. S14G,H), yet dissimilar when traversed in opposing directions (normalized stability = 0.05 ± 0.07, median ± s.e.) despite weak overall heading-direction tuning (Fig. S14I-L). Third, because plane referencing changes only how the grid code is aligned to behavior and not the internal population code itself, activity during unclustered flights should arise from the same toroidal population dynamics as clustered flights, so that a toroidal model fit only to clustered flights should generalize to unclustered flights. Indeed, toroidal latent-variable models^38^ trained only on clustered flights generalized readily to held-out unclustered flights both in the decoded toroidal coordinates (Fig. 4E-G) and in held-out predictions of single-cell activity (Fig. 4H-J). This suggests that the apparent loss of spatial periodicity does not reflect changes in the underlying toroidal population dynamics but may instead arise from combining trajectories across different planes of motion. Thus, the plane-referenced framework helps reconcile our findings with prior reports that did not detect globally periodic firing patterns of grid cells during movement in three-dimensional environments, where neural activity was analyzed without reference to the animal’s movement geometry^18,19^. Finally, alternative grid models that lack plane referencing, including columnar, hexagonal close-packed, and face-centered cubic lattices^18,19,39–41^ (Fig. S15A-E), one-dimensional periodic time or distance codes^42,43^ (Fig. S15A,F,G), and a Lennard-Jones local-order model without global periodicity^18^ (Fig. S16), could not simultaneously account for both periodicity on clustered flights and its apparent absence when unclustered flights were combined.

Together, these results demonstrate that (i) globally periodic grid cells exist in freely flying bats, (ii) their population activity is consistent with an underlying two-dimensional toroidal manifold, and (iii) mapping that manifold onto the animal’s momentary plane of motion accounts for grid-cell activity across both clustered and unclustered flights and provides a parsimonious solution for how a two-dimensional neural code can support navigation in three-dimensional space.

## Discussion

Our findings show that grid cells retain periodic spatial firing patterns during three-dimensional movement and provide evidence for how the interplay between population dynamics and behavioral specialization gives rise to these activity patterns. At the single-cell level, prior studies in bats^18^ and rodents^19^ using volumetric analyses of movement through 3D environments reported irregular or locally ordered firing fields without global periodicity. Our results suggest that this apparent loss of periodicity can be reconciled by considering the ethological geometry of movement: bats do not sample 3D space uniformly but often navigate along locally planar trajectories. By allowing bats to express self-selected flight trajectories and analyzing neural activity in the coordinate frame of their natural behavior, we revealed robust periodic grid-cell activity during 3D movement, highlighting the importance of considering not only the environments animals occupy, but also the species-specific behaviors they naturally express within them^44^. At the neuronal ensemble level, continuous attractor network (CAN) models^1,6–9^ have been successful in describing grid cells during movement in one- and two-dimensional environments, where population activity evolves on a low-dimensional toroidal manifold^10–17,27^. Here, we show that the activity of co-modular grid cells in bats navigating in three-dimensional space exhibits topological signatures consistent with an underlying toroidal network organization, in line with experimental observations in rodents moving in one- and two-dimensional environments^10–17^. Moreover, the persistence of this organization across flight and non-flight periods supports the interpretation that the toroidal structure reflects an intrinsic population manifold, rather than a pattern imposed solely by movement through space, as in models where periodicity is constructed from self-motion inputs^45,46^. Together, these results indicate that, despite their distinct mode of navigation, bats likely rely on similar entorhinal circuit principles as terrestrial mammals, supporting a unified, cross-species framework in which grid-cell activity reflects a shared computational mechanism across behavioral dimensions.

The central question therefore shifts from whether grid-cell periodicity and the underlying toroidal population manifold exist when animals move through three-dimensional space, to how a two-dimensional neural representation can support navigation through a three-dimensional world. Our results suggest that this is achieved by aligning grid-cell representations to behaviorally relevant subspaces, consistent with the dynamic grouping of hippocampal ensembles across spatial reference frames^47^. Furthermore, this framework helps reconcile a set of findings that have been difficult to integrate. In rodents, grid-cell firing can follow the geometry of locomotor surfaces that structure the animal’s motion: fields become vertically elongated on pegboards, helical tracks, and lattices^19,39^, align to steeply sloped terrain^36^, and are altered during wall climbing^48^, observations that motivated the hypothesis that the mammalian spatial map is multi-planar rather than fully volumetric^35,37^. In those studies, the plane of movement was imposed by the terrain, leaving open whether planar coding reflects a property of the neural code or of the locomotor substrate. Yet subsequent volumetric studies in rats and flying bats reported irregular^19^ or locally ordered^18^ firing fields without global periodicity, raising the possibility that grid structure breaks down when movement is no longer constrained to a surface. Our findings suggest a different resolution: during unconstrained 3D navigation, grid-cell periodicity and toroidal organization are preserved, but the periodic structure becomes apparent only when interpreted relative to the animal’s self-selected plane of motion. This could also account for the minority of horizontally aligned grid patterns reported in rats exploring a volumetric lattice layer by layer^19^, where the horizontal plane is the dominant plane of motion^49^. Because in bats this plane changes from flight to flight, the mapping cannot be static: the same toroidal manifold must be flexibly re-referenced to each new plane the animal adopts.

In line with this view, natural flight trajectories in bats, both in the laboratory and in the wild, across spatial scales, are well described by planar subspaces embedded within three-dimensional space, offering a natural behavioral substrate onto which grid activity can be mapped. Such a solution is also consistent with a broader principle of biological design: evolution often repurposes existing implementations rather than constructing entirely new ones^50^. In the case of grid cells, repurposing may reduce the need for higher-dimensional representations, which can carry substantial biological and computational costs^51,52^. Indeed, theoretical work has shown that two-dimensional grid modules can efficiently encode higher-dimensional variables through projections onto planar subspaces, without requiring higher-dimensional grids^51^. Our results suggest that in bats such subspaces can be supplied by behavior: a two-dimensional neural sheet may support navigation in three-dimensional environments by aligning to the animal’s own plane of motion, a tight correspondence between neural dynamics and ethological constraints in which the toroidal manifold and the planar structure of movement may reflect complementary solutions to the same navigational problem. More broadly, this framework yields a testable hypothesis for how a common grid-cell architecture could be deployed across the mammalian tree despite substantial differences in navigational strategies (Fig. S17)^31,32,53–55^: by aligning an evolutionarily conserved two-dimensional neural manifold to the ethologically relevant movement subspaces of each species, the brain may support three-dimensional navigation with a shared architecture rather than species-specific higher-dimensional solutions. Because the reference plane in bats changed from flight to flight, this does not require globally planar movement, only movement that is locally organized into planes. Testing this hypothesis will require comparative studies that leverage the diversity of species and locomotor strategies to determine whether conserved toroidal grid-cell architecture is aligned to species-specific movement subspaces^56^.

An intriguing cross-species question for future work is how an animal determines the plane of motion it occupies, and how the toroidal grid-cell manifold is aligned to it. Across insects, fish, and mammals, directional circuits are often described as one-dimensional ring attractors that provide an azimuthal compass^30,57–63^, which in rodents has been shown to rotate with the locomotor surface^64^. However, specifying a planar reference frame in three-dimensional space requires orientation information beyond azimuth. In bats, head-direction signals extend to three-dimensional angular variables^65^, and vestibular or gravity-referenced signals described in rodents and primates provide plausible sources of this information^66–68^. A plane-referenced grid code thus predicts that the dynamics on the toroidal manifold, and consequently the periodic grid-cell activity patterns, should be quantitatively related to multi-axis directional and gravity-referenced signals so as to project movement in 3D space onto the animal’s behaviorally relevant 2D subspace. This further predicts that such underlying computations should persist even when the local reference frame rapidly changes, such as when the animal transitions between different planes.

Finally, these findings highlight the importance of considering the ethological context of natural behavior when assessing neural computations. Neural circuits are shaped by evolution to support species-typical behaviors, and experimental conditions that restrict or distort those behaviors can obscure the computations they implement^44,56,69–72^. This is particularly important for naturalistic spatial behaviors, where the structure of the animal’s self-generated movement can determine whether an underlying neural code is detectable. Here, grid-cell periodicity in three-dimensional environments was revealed only when neural activity was interpreted in the context of natural flight trajectories. Experimental paradigms that minimize external intervention, including human presence, which itself can shape behavior and neural activity in the hippocampal formation^73^, may therefore better expose the behavioral structure for which these circuits evolved. More generally, understanding brain function requires not only precise measurement of neural activity and behavior, but also paradigms that preserve the natural structure of the behaviors these computations evolved to subserve.

## Materials and Methods

### Subjects

Neural recordings were performed in a total of three adult male Egyptian fruit bats (*Rousettus aegyptiacus*; ∼153–168 g body weight). A total of 13 sessions (3-6 sessions *per* bat) were collected, during which neural activity was recorded wirelessly with a Neuropixels 1.0 probe (NP1.0)^75^ during aerial foraging. All animals were housed in a temperature- and humidity-controlled room. Implanted animals were single housed after implant surgery. Housing room lights were maintained on a reverse light cycle (12 h lights on, 19:00–07:00; 12 h lights off, 07:00–19:00). All experiments were performed during their awake hours (dark cycle) at the same time of the day. All experimental procedures were approved by the Institutional Animal Care and Use Committee of the University of California, Berkeley.

### Aerial foraging

All neural recordings during aerial foraging were performed in a large flight enclosure^23^ (5.6 m × 5.2 m × 2.5 m). During training and recording sessions, bats were mildly food-restricted at >85% of their baseline weight. Bats were trained for 7-14 days prior to performing neural recordings through daily (120 min) sessions in which the bats were allowed to freely obtain pureed fruit reward from automated feeders at designated locations in the enclosure. All trainings and recordings were performed without experimenters inside the flight enclosure to avoid confounds due to the presence of humans^73^. Recording sessions lasted between 80 and 150 minutes.

The flight enclosure is acoustically, electrically and radio-frequency-shielded, and equipped with high-precision lighting control. The room ceiling and walls were covered with acoustic foam to minimize acoustic reverberations and to dampen noise from outside the room. The walls and floors were covered with an additional layer of acoustically absorbing black felt to protect the acoustic foam from being damaged, and to provide a perch-able substrate for the bats. A set of 16 motion-capture cameras^23,24^ (Raptor-12HS, Motion Analysis) was used to track the 3D spatial position of the bats with millimeter-resolution. Three reflective markers, asymmetrically arranged on the neural recording headstage on the head of the bat, were tracked by the motion capture cameras at a frame rate of 120 Hz using commercially available software (Cortex-64; Motion Analysis). Two automated feeders, which dispense a pureed fruit reward, were positioned 1.8 m apart anchored to the wall at one end of the room. Two feeders were used to encourage diversity in flight paths by providing bats with multiple options to forage from, while still maintaining sufficient repeated trials of each flight path. Similar to in previous studies^23,24^, reward was triggered by an infrared beam break sensor that was interrupted when a bat landed on the feeding platform. To receive reward, bats were required to fly a minimum of 3 meters away from the feeder each time before retriggering the beam break. Each feeder was independently controlled by an Arduino (Uno Rev3) and Adafruit Motorshield (1438; Adafruit) which interfaced with a computer outside of the experimental room. Reward probability (0.2 – 0.8) and amount (0.1-0.2 mL) were adjusted by the experimenter to fine tune the bat’s behavior. All recording sessions were conducted under uniform illumination (illuminance of 5 lux).

All recording systems (position tracking and neural recordings) were synchronized by ground truth periodic clock pulses generated by a Master-9 device (A.M.P.I.) that served as the central clock for the experiments. On-board accelerometer data were acquired at 500 Hz by the neural recording headstage in parallel with the neural data.

### Surgery

Probe implants were performed in two stages, separated by 7-10 days: (i) implantation of the training cone, and (ii) insertion of Neuropixels 1.0 probes. The surgical procedure was performed as previously described in detail^25^.

#### Implantation of the cone

Anesthesia was induced using an injectable cocktail of ketamine, dexmedetomidine and midazolam. The bat was then placed on a stereotaxic apparatus (Model 942; Kopf) and provided with a continuous supply of oxygen. Anesthesia was maintained via injections of a cocktail of dexmedetomidine, midazolam, and fentanyl (∼one dose per hour). Anesthesia depth was monitored continuously by the bat’s breathing rate and by a toe pinch reaction test. Body temperature was continuously monitored using a rectal temperature probe and maintained at approximately 35°C. Each Neuropixels probe was grounded using a stainless-steel wire (0.008 in. coated; A-M Systems) soldered to a ground screw (19010-00; FST), which was inserted into the frontal plane of the skull. Four shorter bone screws (M1.59 mm stainless steel) were inserted into the skull to further strengthen the attachment of the implant. A 1 mm circular craniotomy was made at the probe insertion point (at approximately 0.5 mm anterior to the transverse sinus and 4.9 – 5.7 mm lateral to the midline). The craniotomy was sealed with a biocompatible elastomer (Kwik-Sil; World Precision Instruments), and a custom training cone was cemented. At the end of the surgery, reversal agents were injected to counteract the dexmedetomidine and midazolam. An oral analgesic (Metacam; Boehringer Ingelheim) was administered after the bat had fully awoken from the anesthesia. Analgesics (3 days) and antibiotics (7 days) were provided daily post-op until full recovery. Behavioral training was resumed after full recovery from surgery (about four days). Implant weight was gradually increased during behavioral training to allow the bat to adapt to the full implant.

#### Insertion of Neuropixels 1.0 probes

Prior to probe insertion, each Neuropixels 1.0 probe (NP1.0) was sharpened at a 20-30° angle for 15 minutes using a Microgrinder (EG-45; Narishige), and a single stainless-steel wire (0.008 in. coated; A-M Systems) was soldered to the ground and reference of the probe. The probe insertion procedure follows the same general surgical practice as described above and previously^25^. After the bats were anesthetized and placed in a stereotaxic apparatus, the probe shank was coated with fluorescent dye (CM-DiI; Invitrogen C7001) and inserted into the target craniotomy at a rate of ∼10-20 µm/sec at a pre-calculated angle. The initial insertion angle was calculated for the exact craniotomy location of each bat individually (between 20° to 40° azimuthal angle from the sagittal plane, and 15° to 25° pitch angle from the dorsal-ventral axis, such that the probe travels from posterior to anterior, medial to lateral). The probe was inserted to the maximum depth at which visible bending of the probe shank was observed (skull depth). If necessary, the probe was retracted and re-inserted following the above procedure with incremental changes in the insertion angle (10° azimuth and 2° pitch per step) until a desired skull depth of 6000-6500 µm was measured. On the final insertion, the probe was retracted by 150-300 µm to prevent contact with the skull during recordings. The probe was then cemented in place using dental acrylic and the ground wire of the probe was connected to the pre-existing ground screw. A new 3D-printed protective cone was positioned and cemented to the skull and the bat was awoken using reversal agents to counteract the dexmedetomidine and midazolam.

### Electrophysiology data acquisition, pre-processing, and spike sorting

Recordings began one day after probe insertion (5 hours post-insertion for one bat) and were performed using a SpikeGadgets wireless NP1.0 headstage as previously described^25^. Channels were selected to record from dorsal MEC and were provisionally determined by a visible increase in spiking activity, typically around 4000 µm depth (from probe tip) and subsequently verified histologically. Electrical signals (referenced to the ground screw) in the spike band (600 – 6000 Hz) and LFP band (0.5-200 Hz) were amplified 1000x – 1500x and 250x respectively. Drift correction and spike sorting were performed automatically using Kilosort4^76^. All units labelled by Kilosort4 as *good* were then manually curated in Phy^77^. Rastermap^74^ visualization of population activity was computed using gaussian-smoothed (σ = 50 ms), z-scored firing rates on a 600 s long segment of a session with the following parameters: 10 ms time bins, ‘n_clusters’ = 96, ‘n_PCs’ = 128, ‘time_lag_window’ = 50, and ‘locality’ = 0.3.

### Histology

At the end of the planned experiment sessions, bats were given a lethal overdose of sodium pentobarbital and perfused transcardially (200 ml phosphate buffered saline, 0.025M, pH = 7.4; 200 ml of fixative, 3.7% Formaldehyde in phosphate buffered saline). The probe implant was carefully removed, and the brain was dissected and stored in the fixative solution (3.7% Formaldehyde in phosphate buffered saline) for 1-2 days, then in a 30% sucrose solution in phosphate buffered saline overnight for cryoprotection. A microtome (HM450, Thermo Fisher Scientific) with a freezing stage was used to cut 50-µm sagittal sections. Slices containing the medial entorhinal cortex were stained for DAPI and coverslipped with an aqueous mounting medium (ProLong Gold Antifade Mountant with DNA stain DAPI, Thermo Fisher Scientific). Fluorescent images of sagittal sections were acquired using an Axioscan Slide Scanner (Zeiss) and Neuropixels probe tracks were visualized and localized by CM-DiI fluorescence.

### Processing of position tracking data

Positional data recorded by the marker-based motion capture system (acquired at 120 Hz) were preprocessed^24^, to obtain continuous and smooth 3D positional data. Flight trials were segmented based on a velocity threshold of 1 m s^−1^ and 3D spatial trajectories during flight were clustered using hierarchical clustering^24,25^. In brief, flight trajectories were downsampled in time for clustering by a factor of 12, and clustered using agglomerative hierarchical clustering via Fréchet distance^25,78^. The linkage distance was selected to be the highest threshold at which the silhouette score increased from the previous highest silhouette score by more than 0.05. Flight segmentation boundaries were refined by time-lagged cross-correlation alignment of the on-board accelerometer signal, to match the exact time of the first wingbeat. Mean flight duration was 3.42 ± 1.21 s (mean ± s.d.). Flights with unfillable missing tracking near take-off or landing were excluded, and clusters with at least 3 well-tracked flights were defined as clustered flight paths (3 to 35 flights per flight cluster). All flight clusters with fewer than 3 flights were pooled together into the unclustered flights (6 – 50 unclustered flights per session). Clustered flights have a mean duration of 3.08 ± 0.80 s (mean ± s.d.), with shortest clustered flight duration being 1.11 s. The distribution of clustered flight durations was bimodal, with straight flights having approximately half of the duration of loop flights (straight flights, 1.65 ± 0.24 s; loop flights, 3.47 ± 0.30 s; mean ± s.d.).

### Defining stable multifield neurons

Stable multifield neurons were defined as those with (i) a minimum of 75 spikes on at least one clustered flight path within its 2D rate map, (ii) a minimum of 4 firing fields on at least one clustered flight path, and (iii) an odd flight vs even flight spatial rate map Pearson correlation of at least 0.3. Of the 2,353 total recorded cells, 1,253 cells met the spike-count and field-count criteria and were classified as multifield; of these, 1,102 additionally met the odd/even stability criterion and were classified as stable multifield (Fig. S3B shows the odd/even correlation distribution for all 1,253 multifield cells). The stability criteria of odd vs even spatial rate maps were constructed by splitting clustered flights into odd and even trials. Odd trials from all clustered flights were combined into the odd group, and similarly for the even group. 2D spatial rate maps were then computed (see ‘Rate map estimation’) for the odd group and even group separately, using a bin size of 15 cm, gaussian kernel of 1.2 bins, and minimum occupancy threshold of 50 ms. To estimate the number of firing fields on a clustered flight path, we first computed the 2D spatial rate map on each clustered flight path as done above (see ‘Rate map estimation’), then applied an additional Gaussian smoothing step (σ = 1.2 bins) for peak detection only, and performed local maxima detection (using skimage function ‘peak_local_max’) of all peaks exceeding the mean rate map firing rate with minimum inter-peak Chebyshev distance of 2 bins. For these criteria, only clustered flight paths with at least 5 flights were considered (the odd/even split pooled flights from clustered paths with at least 6 flights). Subsequent analysis of each multifield neuron (see ‘Quantification of 1D periodicity’) was performed only on the subset of flight clusters in which it had sufficient spikes and multiple firing fields.

### Quantification of 1D periodicity

To quantify the periodic firing of grid cell responses on clustered flight paths, spikes were binned into 10 ms time bins and neural activity was trial aligned to the time of take-off. Take-off time estimates were refined by finding the time-lag that minimized the mean squared error between each trial’s position trace and the cluster mean. Trial averaged firing rates were computed from the binned spike counts and smoothed with a 20 ms gaussian kernel. The autocorrelation of the trial-aligned firing rates was computed up to max lag equal to 50% of the trial length. Power spectral density (PSD) was computed from the trial-averaged firing rates using Welch’s method with window size equal to 80% of the trial length with 90% overlap. For visualization in Fig. 1J, a window size of 50% of the trial length was used. Cells were classified as significantly periodic if, on at least one flight path, there existed a peak (minimum prominence of 20% of the maximum) between 1 Hz and 5 Hz that exceeded the 95^th^ percentile of shuffle (corresponding to the possible observed range of detectable grid spacings given the size of the flight enclosure, approximately 4 to 20 firing fields along a flight path). The 95^th^ percentile PSD threshold was Bonferroni-corrected for the number of flight paths each cell was evaluated on. Shuffled PSD distribution (*n* = 1,000 shuffles) was computed by applying a random circular shift, between 0 s and the trial duration, to the spike train of each flight trial independently, thereby breaking the correspondence between spikes and position along flight paths while preserving the intrinsic spike train properties.

### Power spectral density clustering

Significantly periodic grid cells (as defined in ‘Quantification of 1D periodicity’) were pooled across sessions and clustered based on their z-scored power spectral density (PSD), in the 1 – 5 Hz band, along a representative flight cluster using agglomerative clustering with average linkage and correlation distances at threshold *ρ* = 0.5. For each cell, the representative flight cluster was chosen as the cluster with maximal fraction of total spectral power between 1 Hz and 5 Hz contained within a ±0.3 Hz window around the largest PSD peak in that range, and a minimum flight duration of 3 s. For visualization, a minimum cluster size threshold of 50 cells was used. The average grid spacing of each resulting cluster was estimated from the mean PSD peak frequency, and clusters with peak frequency below 1.2 Hz were excluded. Autocorrelation heatmaps were then constructed by stacking each autocorrelation cluster sorted by their average grid spacing. For per-bat autocorrelation heatmaps, autocorrelation clusters were sorted by their average recording depth, and split by bat identity, and a minimum cluster size of 15 cells was used for visualization.

## Grid module ensemble detection

Detection of co-recorded grid module ensembles was performed using a spike-time-correlation-based method as proposed by Hermansen et al.^14^. In brief, we estimated the firing rate, ***s****_i_*(*t*), of each neuron by binning the spike times into 10 ms time bins and smoothing with a σ = 20 ms gaussian kernel. Low firing rate cells (threshold between 0.25 – 1 Hz, mean session rate) were excluded. We then computed the cross-correlation between each pair of co-recorded neurons across the full recording session (spanning all behavioral states, including flight and non-flight bouts):

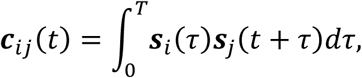

where T denotes the length of the full recording session. Next, we computed the inverse, normalized cross-correlation matrix as:

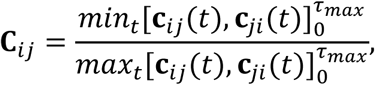

where τmax was chosen per recording between 100 – 200 ms (P1, P2, P3, R1: τmax = 100 ms, Q1: τmax = 200 ms). Finally, we computed the correlation distance between each row of **C**^2^ to identify ensembles of neurons that have stable within-module relations, as grid cells should be consistently in phase or out of phase with each other. We then performed agglomerative clustering with average linkage using a distance threshold of *ρ* (P1: *ρ* = 0.375, P2: *ρ* = 0.475, P3: *ρ* = 0.475, Q1: *ρ* = 0.45, R1: *ρ* = 0.3). Ensemble detection code was based on implementations in the code repository by Hermansen et al.^79^. For grid module identification, all recorded time bins, including both flight and rest bouts were used for computing the correlation distance matrix. For flight versus non-flight comparisons, distance matrices were computed using only in-flight time bins or non-flight time bins, respectively, and the cells with no cross-correlation overlap in spikes within τ_max_ were excluded from the visualization. For visualization, firing rates were smoothed with a σ = 30 ms gaussian kernel, and τ_max_ = 100 ms.

Pairwise cross-correlograms for co-modular cell pairs were computed following procedures adapted from Trettel et al.^11^. In brief, cross-correlograms of the mean-centered, smoothed firing rates (σ = 20 ms) were averaged across flight and non-flight bouts respectively. For visualization, each row of the correlogram was mean-centered within a ±1 s lag window, and ordered by zero-lag correlation value according to flight correlogram, and the same ordering was applied to the non-flight correlogram.

### Preprocessing of population activity

Preprocessing of population activity for persistent cohomology was adapted from procedures previously used in rodent studies^13,14^. Firing rates were computed by binning spikes into 10 ms time bins and smoothed with a 20 ms gaussian kernel to obtain population activity vectors (firing rate of each neuron in an ensemble). The population activity was temporally downsampled by a factor of 2 for in-flight time bins. Non-flight bouts were more heavily downsampled in time to reduce computational burden due to the large amount of time bats spend resting in a session (P1: 15, P2: 20, P3: 20, Q1: 20, R1: 10). The population activity vectors were sorted by total activity, and the top fraction of time bins was selected to remove time bins with low or zero activity (Flight bouts, P1: 0.85, P2: 0.65, P3: 0.8, Q1: 0.6, R1: 0.85; Non-flight bouts, P1: 0.65, P2: 0.75, P3: 0.7, Q1: 0.45, R1: 0.45). The firing rate of each neuron was then z-scored, and the population activity vectors were projected into a 6D PCA subspace and scaled to unit variance, as done previously^13,14^.

Next, we performed a radial downsampling algorithm as proposed by Hermansen et al.^14^ to further reduce the size of the point cloud while preserving the spread of the data. In brief, the Euclidean distance of the point cloud to the point with maximal total magnitude (initial landmark point) was computed. Points within *ɛ* of the most recent landmark were then discarded and the remaining point with the smallest maximum distance to all previously chosen landmarks was picked as the next landmark point. This process was repeated until no non-landmark points remained (flight bouts: P1, *ɛ* = 0.25; P2, *ɛ* = 0.05; P3, *ɛ* = 0.05; Q1, *ɛ* = 0.3; R1, *ɛ* = 0.01; non-flight bouts: P1, *ɛ* = 0.4; P2, *ɛ* = 0.15; P3, *ɛ* = 0.15; Q1, *ɛ* = 0.35; R1, *ɛ* = 0.05). Radial down sampling preserves the point cloud shape but is prone to outliers, thus a second density weighted down sampling procedure was performed as described previously^13,14^ to reduce outliers and yield the final point cloud with desired size. In brief, the neighborhood strength of each point to its k nearest neighbors was defined as 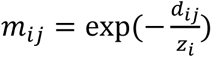, where *d_ji_* is the cosine distance between points i and j, and the normalization factor *z_i_* was chosen such that *m_ij_* sums up to *log*_2_(*k*), with k = 1000. The symmetrical neighborhood strength is defined as *M_ij_* = *m_ij_* + *m_ji_* − *m_ij_* · *m_ji_*. This down sampling procedure was iteratively repeated until the final N points were chosen by picking at each step the point with largest summed neighborhood strength and subtracting that point’s pairwise strengths from the summed strengths of the remaining points (Flight bouts, P1: 1800, P2: 1200, P3: 1200, Q1: 1500, R1: 1300; Non-flight bouts, P1: 1650, P2: 1350, P3: 1650, Q1: 1500, R1: 1350). Finally, the negative logarithm of the downsampled point cloud neighborhood strengths was computed to yield a fuzzy distance matrix (k = 800), which is then provided to the Ripser^80,81^ implementation for persistent cohomology computations.

### Persistent cohomology

To quantify the topological features in co-modular grid cell population activity (see ‘Preprocessing of population activity’), we used persistent cohomology, a topological data analysis technique previously used successfully for quantifying head direction ring topology^29,30,82^ and grid cell toroidal topology^13,14^. Persistent cohomology is closely related to persistent homology, yields identical persistence barcodes, but is more computationally efficient and enables cohomological decoding from computed cocycles^83^. The detailed theoretical framework of persistent cohomology is as described elsewhere^13,14,83^. In brief, a ball of radius r is placed at each point in the point cloud. As r is increased from zero, overlapping spheres merge, building up a sequence of connected shapes (a filtration) in which topological features such as connected components (0-D holes), circular features (1-D holes), and cavities (2-D holes) emerge at some radius, and are eventually filled. The range of radii at which a topological feature is present is denoted as bars in the persistence barcode, with longer bars indicating more robust topological features. Persistent cohomology computations were performed with Ripser^80,81^. Persistent cohomology was applied to grid cell population activity during flight bouts (including all clustered and unclustered flights), and separately for non-flight bouts. Identified modules were labeled by letters indicating the bat identity (P, Q, R) and numbers (e.g., P1, P2, P3) indicating multiple modules from the same animal.

To confirm if the observed lifetimes of topological features exceed what is expected by chance, a random circular shift in time, from 0 to the total flight duration or rest duration, was applied to the activity of each neuron independently, and the topological lifetimes of each shuffle (*n* = 500) were computed using persistent cohomology following the same procedure as described above. To verify that the observed toroidal signature is above chance, we compared the observed barcode against toroidal signatures obtained in shuffled point clouds. Significance criteria of H0, H1, and H2 were set to the 95^th^ percentile of the longest finite H0 bar, the second longest H1 bar, and the longest H2 bar, respectively.

### Cohomological decoding of toroidal coordinates

Toroidal coordinates were decoded from the internal population dynamics via cohomological decoding introduced by de Silva et al.^83^, which has been applied previously for the study of head direction and grid cell population activity^13,14,29^. The implementation used here, described in brief below, is based on the published methods and code from Gardner et al.^13^ and Hermansen et al.^14,79^. Persistent cohomology was computed on the downsampled point cloud of grid cell activity (see ‘Preprocessing of population activity’ and ‘Persistent cohomology’). Circular-valued coordinates were then decoded for each of the two longest-lived H1 bars from their cocycle representatives by lifting to integer coefficients and smoothing at a filtration scale τ = b + 0.99·(d − b), where b and d denote the birth and death time of the second longest H1 bar. To improve robustness to non-uniform sampling of the underlying manifold, smoothing was performed under successively higher L^p^ norms (p = 2, 4, 6, 10, 20) and finally under the L^∞^ norm, following procedures established in Paik & Park^84^. Interpolation of toroidal coordinates to time points beyond those of the downsampled point cloud, both within and across behavioral states, was done by weighting the coordinates of the downsampled points by their PCA-projected population activity, yielding a distribution on the torus for each principal component. Toroidal coordinates at a query time bin were then computed as the center of mass of these distributions, weighted by the PCA-projected population activity at the query time bin.

### Rate map estimation

#### Spatial rate map

To compute 2D spatial rate maps, spike and bat positions were binned into 10 cm × 10 cm bins, and bins with less than 30 ms occupancy were excluded. The binned spike counts and binned occupancy were then smoothed with a gaussian kernel (σ = 1.5 bins). Spatial rate maps were then estimated by dividing the smoothed binned spike counts by the smoothed binned occupancy, and bins without a neighboring bin (or itself) with at least 30 ms raw occupancy were excluded. 3D rate maps, unless stated otherwise, were estimated using a similar procedure as 2D rate maps using 15 cm × 15 cm × 15 cm bins, σ = 1.2 voxels smoothing, and minimum occupancy of 50 ms. For visualization only, 3D rate maps were interpolated into a finer grid (half the voxel size within layer and 5 cm between layers). No interpolation was applied for visualization of 2D rate maps.

#### Toroidal rate map

Toroidal rate maps were estimated from the spike counts and decoded toroidal coordinates (see ‘Cohomological decoding of toroidal coordinates’) per 10 ms time bin. The full toroidal phase space was divided into 20° × 20° bins, and toroidal phase occupancy was computed by binning the decoded toroidal coordinates. Toroidal phase spike counts were tabulated by counting the number of spikes that occurred in each toroidal phase bin. Before smoothing, each row in the toroidal phase occupancy and spike count maps was rolled horizontally by half its vertical index to account for the 60° angle between the toroidal axes. The sheared maps were then tiled into a 3 × 3 grid, smoothed with a gaussian kernel (σ = 1.5 bins), and cropped to the central tile (following similar procedures used in rodents^13,14^). The toroidal rate map was then computed as done for spatial rate maps by dividing the smoothed spike counts by the smoothed occupancy maps, and bins with smoothed occupancy below 10 ms were excluded. For visualization only, the axes of the toroidal rate map were sheared by 15°, and NaN bins were iteratively in-filled by the mean of their neighboring non-NaN bins starting from the bin with the most-neighbors (respecting the 60° toroidal axes angle as done above).

Pairwise similarity between cells’ spatial rate maps (computed per cluster in 2D, then averaged across clusters) or toroidal rate maps is measured with the Pearson correlation between flattened rate maps on shared occupancy bins. Correlation between pairwise spatial similarity and pairwise toroidal similarity was quantified by Pearson correlation in comparison to a shuffle distribution generated by randomly permuting the cell identities within session (*n* = 1,000 shuffles).

## Toroidal phase map estimation

Toroidal phase maps were computed by binning the toroidal coordinates at each bat position (projected onto the flight plane, or top-down projected) into 10 cm × 10 cm bins and taking the circular mean of the toroidal coordinate within each bin. The resulting phase maps were smoothed by applying a gaussian kernel (σ1 = 0.6 bins) to the sine and cosine projections of the phase and recombining them with arctan2 to obtain angular phase maps (used for ‘Fitting of plane waves’). A minimum occupancy threshold of 10 ms was applied to the smoothed occupancy. Cosine-projected phase maps in Fig. 3G, Fig. 4F, and Fig. S8E, used an additional gaussian smoothing step (σ2 = 0.8 bins) after cosine-projection for visualization. Phase map similarity across conditions (Fig. 4G and Fig. S8E,F) was measured by the mean vector length (MVL) of the angular difference between phase maps.

### Stability of toroidal tuning

To assess the stability of grid cell tuning to the underlying toroidal manifold across behavioral states, a separate toroidal parameterization was computed using flight bouts and non-flight bouts (see ‘Persistent cohomology’ and ‘Cohomological decoding of toroidal coordinates’). Toroidal rate maps during flight bouts were computed from the flight bout toroidal parameterization, and rate maps during non-flight bouts were computed from either (i) the same flight bout toroidal parameterization, or (ii) a separate parameterization computed from non-flight time bins. When using the same flight-bout toroidal parameterization, the similarity between toroidal tuning during flight and non-flight is measured by the Pearson correlation between the toroidal rate maps. With separate toroidal parameterizations for flight and non-flight, correspondence between (*ϕ1*, *ϕ2*) obtained during flight and during non-flight is not guaranteed as the decoded coordinates may correspond to any two of the three possible axes of a hexagonal lattice and may also differ in choice of origin and orientation. Following previously described procedures^14^, we smoothed the decoded flight-time toroidal coordinates obtained from each parameterization in time with a gaussian kernel (σ = 20 ms), then tested possible (*ϕ_1_*, *ϕ_2_*) correspondences and picked the transformation that yielded the minimum angular error between flight and non-flight parameterizations. For each transformation, phase shifts were analytically calculated from the circular mean of the residual angular differences. For both (i) and (ii), the module-level stability was computed as the mean correlation across co-modular cells. Significance was assessed with random permutations of cell identity on non-flight bouts within session (*n* = 1,000).

### Analysis of flight geometry

#### Flights in the wild

We analyzed the geometry of Egyptian fruit bat flight during large-scale navigation in the wild using a publicly available GPS-tracking dataset from Egert-Berg et al.^33,34^, which tracked free-ranging bats commuting in the wild between roosts and distant foraging sites. Position traces were preprocessed using a 15-sample median filter followed by a Gaussian filter (σ = 2 samples). Missing gaps of ≤3 samples were linearly interpolated. Initial flight segmentation was based on a velocity threshold of 10 km h⁻¹, after which flight boundaries were refined by extending each segment (up to 25% of the initial segment length) to the nearest sample with altitude <5 m. Flights were then filtered for commute trajectories with total distance traveled ≥10 km and straightness index^31,33^ ≥ 0.7 where straightness index was defined as the ratio of the shortest path between flight endpoints to the actual path length. The cruise phase of each flight was defined as the portion of the trajectory between the first and last samples exceeding 75% of the maximum flight altitude (defined as the 99^th^ percentile of height for robustness to noise). Ascent and descent phases were defined as the trajectory segments preceding and following the cruise phase, respectively, with a minimum of 1 km horizontal travel distance. Flights with ascent or descent phases spanning <50 m altitude change, or cruise phases shorter than 5 km, were excluded from further analysis. Each flight phase was fit independently using piecewise planes from K = 1 to 5 planes. For each K, the trajectory was partitioned into K contiguous segments, with minimum length of 10 samples for wild data and 20 samples for laboratory data, that minimized the total L1 residual (perpendicular distances of samples to their corresponding plane). Piecewise variance explained was defined as:

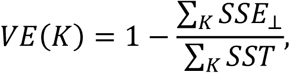

where *SSE*_⊥_ denotes the sum of squared residuals of each segment from its PCA plane, and SST is the total sum of squares of each segment from its centroid. For K=1, this reduces to the standard variance explained by a single best-fit plane. A flight phase is defined to be well-fit by K planes if the piecewise variance explained is greater than 0.98.

#### Flights in the laboratory

Following preprocessing of foraging flights collected in the laboratory (see ‘Processing of position tracking data’), the same piecewise plane fitting procedure described above was applied to each individual flight trial, including both clustered and unclustered flights. Flights were classified according to the minimum number of planes required to achieve piecewise variance explained > 0.98. Plane of motion was defined by the first two principal components, **u** and **v**, of the positions in a flight cluster (or flight trajectory). The sign of **u** was chosen such that **u** points from the take-off position in the direction of the farthest position from take-off, and sign of **v** was chosen such that **u** × **v** has a positive z component. The angle of a plane was given by θ = arccos(|**n̂** ⋅ **ẑ|**), where **n̂** is the unit normal to the plane and **ẑ** is the unit vertical axis. The inter-plane-angle between planes i and j was defined as θij = arccos(|**n̂**i ⋅ **n̂**j|), which is the minimum angle between the two planes under azimuthal rotation. Because near-straight flights have less stable roll estimates (rotations about PC1), quantification of plane angles was done using loop flights with well-defined normal vectors.

### Fitting of plane waves

To fit the toroidal phase maps on the flight plane with plane waves, we performed a grid search over all plane wave angles (0 to 179° in steps of 1°) and grid spacings (125 geometrically spaced steps between 0.6 and 2.5 m). Optimal plane wave phase was computed analytically for a given angle and spacing. As *ϕ1* and *ϕ2* are offset by 60° or 120° for hexagonal grids, we fit the *ϕ1* and *ϕ2* phase patterns simultaneously, with a fixed angular offset between the plane waves for each. The mean vector length (MVL) of the angular error distribution was used as the fitting metric (higher MVL indicates better fit and is bounded between 0 and 1). For each module, a shared grid spacing was chosen by picking the spacing that gave the highest average MVL across clustered flights. Since *ϕ1* and *ϕ2* could correspond to plane waves either 60° or 120° apart, the fitting procedure was repeated for both angular offsets and the one that yielded the highest average MVL score on clustered flights for each module was chosen. Significance of the plane wave fit for each module was assessed by the average MVL at the shared spacing relative to the 95^th^ percentile of the shuffle distribution (*n* = 1,000 shuffles). Shuffle distribution was constructed by applying a random circular time shift, between 0 s and the trial duration, to each dimension of the toroidal coordinates for each flight trial independently.

### Candidate grid cell selection

Candidate grid cells were identified from the inter-spike-distance distribution of each unit. For each unit, the pairwise 3D Euclidean distances between every within-flight spike pair from all trials of all clustered flights were binned into a 1D histogram (99 bins from 0 to 5 m). The distribution was smoothed with a Gaussian kernel (σ = 1.5 bins) and normalized by the maximum height at distances >0.5 m. Peaks were detected with the SciPy function ‘find_peaks’, using a minimum prominence of 0.2 and a minimum separation of 5 bins. A unit was included if it satisfied: (i) ≥ 150 spikes emitted during clustered flights; (ii) at least two periodic peaks ≥0.5 m, with at least one ≤ 2 m, and (iii) a central peak <0.5 m corresponding to within-field distances.

### Fitting of hexagonal grids on planes of motion

To fit grid patterns on the plane of motion to grid cell activity, the observed spikes and bat positions were projected onto the flight plane. We performed an exhaustive grid search of ∼2.7 million candidates spanning the full landscape of grid spacing (125 geometrically spaced steps from 0.6 m to 2.5 m), orientation (0° to 59° in 1° steps), and xy phase (19 × 19 = 361 total phase bins tiling the full phase space). For each grid parameter combination, a candidate rate map was constructed with hexagonally arranged firing fields with the given grid spacing, orientation, and phase. Grid fields were modeled as Gaussian profiles with *σ* = 0.125 · *s*, where *s* is the grid spacing. Each flight cluster was fit independently using the Poisson log-likelihood of the observed spike positions as given by:

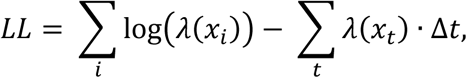

where *λ*(*x_i_*) is the firing rate of the candidate grid pattern at the position of spike *i*, *λ*(*x_t_*) is the firing rate at the bat position at time t, and Δ*t* is the time bin size (10 ms). Positions and spikes were binned in 0.1 m spatial bins, and *λ* was evaluated at the bin center. The amplitude of *λ* was fit so the expected number of spikes under the model matched the observed spike counts. The shuffle distribution is constructed by applying random circular shift to each trial of a flight cluster, between 1 and N – 1 time bins, where N is the number of time bins in each trial, and performing the same grid fit procedure as described above. The spacing of each cell was estimated by summing across flight clusters the maximum log-likelihood at each grid spacing, and picking the shared grid spacing with the highest z-scored total log-likelihood relative to that of shuffles. Each shuffle selects its own shared grid spacing, with z-scores evaluated relative to the remaining shuffles not including itself. A grid fit was considered significant if its total log-likelihood exceeded the 95th percentile of the shuffle distribution (*n* = 1,000 shuffles).

Grid fitting on planes rotated from the plane of best fit was performed by projecting the spike positions onto each rotated plane of motion and applying a similar fitting procedure as above including on shuffled data (*n* = 1,000 shuffles). Planes were rotated away from the best-fit plane, from -40° to +40° in 10° steps at each of 6 azimuth steps from 0° to 150° in 30° steps for a total of 49 planes including the best-fit plane. Grid spacing was selected independently for each flight cluster on each rotated plane. For each cell, fitting on rotated planes was restricted to flight clusters on which it had a significant grid fit on the cluster’s plane of best fit (*n* = 283 cells with at least one valid cluster on which its fit exceeded the 95^th^ percentile of shuffles). To verify the effect was not due to selection of significant grid cells based on the best fit plane, the analysis was repeated using all flight clusters where the cell was significant on any of the rotated planes, Bonferroni-corrected for multiple comparisons across the 49 planes (n = 273 cells with at least one valid cluster). Non-grid multifield cells (*n* = 58 cells) were defined as cells with no significant fit on any rotated planes (Bonferroni-corrected). Simulated grid and non-grid multifield cells were generated on the plane of motion of each flight cluster (n = 360 grid and n = 360 non-grid cells). Grid cells were simulated on the plane of best fit with random spacing (0.8 m to 1.5 m), field size (σ = 0.1 × spacing to σ = 0.15 × spacing), orientation (0° to 60°), and phases. Firing rates were sampled from the in-flight firing-rate distribution of recorded cells. Non-grid multifield cells were simulated with matching field count within 3σ distance of visited positions by the bat to each corresponding simulated grid cell, and a minimum separation between fields of 0.3 m. To ensure simulated non-grid cells were not grid-like by chance, each was simulated 50 times and the least grid-like, measured by rate map correlation with a hexagonal lattice, was selected. The same grid fitting on rotated planes procedure was then performed on the simulated grid and non-grid multifield cells, with n = 100 shuffles. To quantify how lattice fit quality changed on rotated planes, we took the average change in z-scored log-likelihood for each cell at each rotation angle θ, pooling across azimuths and directions, and estimated the slope, β, of this change as a function of θ. Significance was assessed using a one-sample two-sided Wilcoxon signed-rank test comparing the distribution of β across cells with zero. For simulated data, β was computed using the relative z-score defined as the ratio of the z-score on each rotated plane to that on the best-fit plane. Significance for simulated data was assessed by comparing the distribution of paired differences in β (simulated grid cell minus its matched non-grid multifield cell) with zero using a one-sample Wilcoxon signed-rank test.

### Alignment of plane-referenced grids

To compute the alignment between grid patterns on different flight clusters, we estimated the orientation and phase of the grid on each flight cluster under a shared spacing following a similar grid search procedure as before (see ‘Fitting of hexagonal grids on planes of motion’), spanning the grid parameter space across spacing (75 steps, 0.6 m to 2.5 m), orientation (60 steps, 0° to 59°), and phase (14 x 14 = 196 steps). The shared spacing was chosen to maximize the pooled Poisson log-likelihood across all flight clusters under a single shared firing rate amplitude. We then performed a rotation and translation on each flight cluster equal to the fitted orientation and phase of its respective grid, bringing all flight clusters into a common plane-referenced coordinate system. As alignment requires reliable estimation of grid parameters, which improves with more spikes and longer flights, we selected cells with ≥300 spikes on sessions with at least three long (≥8m) flight clusters, yielding a cohort of *n* = 136 cells. The combined coverage of multiple flight clusters allowed a gridness score to be computed from the autocorrelation of the aligned rate map, similar to that used for grid cells in 2D^4,85^. In brief, aligned rate maps were computed from the aligned spikes and flight positions as described in ‘Rate map estimation’ using Gaussian smoothing of σ = 0.8 bins. The autocorrelation of the aligned rate map was computed, and an annulus from 0.5 × s to 1.3 × s was extracted, where s is the estimated shared spacing. The rotational autocorrelation of the annulus was computed in 2° steps, and the gridness score was defined as the minimum correlation at 60° and 120° minus the maximum correlation at 30°, 90°, and 150°. Room-referenced gridness was computed identically, including the same shared-spacing estimation and gridness computation, except that the alignment step was omitted. Shuffled data (*n* = 1,000 shuffles) were generated by circular shifting the spikes on each flight trial by a random amount between 0.1 × T and 0.9 × T, where T is the trial duration, and all grid fitting, spacing estimation, and grid alignment (for aligned plane-referenced gridness) steps were applied identically to each shuffle. For each cell, gridness was considered significant if it exceeded the 95^th^ percentile of the corresponding shuffle distribution (aligned plane-referenced gridness against aligned shuffles, room-referenced gridness against room-referenced shuffles). For each cell and coordinate frame, gridness was also expressed as a z-score relative to the corresponding shuffle distribution. Per-cell differences in z-scored gridness between frames were compared with a two-sided Wilcoxon signed-rank test, and the numbers of cells with significant gridness in the two frames were compared with a two-sided McNemar exact test (40 cells significant in plane-referenced coordinates only, 3 in room-referenced coordinates only, 6 in both, and 87 in neither). Binomial tests comparing the fraction of cells with significant gridness (relative to 5% chance level) were one-sided.

### Grid cell stability and periodicity on mixtures of flight planes

To quantify the effect of pooling together flights on different planes on the observable grid cell periodicity, we pooled together the flight clusters in all combinations from K = 1 to K = 5 clusters, as well as an ‘All’ cluster containing all flights including unclustered flights. This analysis was performed on grid cells with significant grid fits (on best-fit plane) from sessions with at least 5 distinct clustered flights (*n* = 197 cells). We computed a normalized prominence score, defined below, as a measure of apparent lattice structure for grid cells with significant grid fits. For each combination of K flight clusters, we first computed the ISD by binning all cross-flight inter-spike-distances (excluding within flight pairs) into 0.05 m bins, with each spike pair (i, j) weighted by 1⁄(*T_i_T_j_*), where Ti and Tj are the number of trials in their respective clusters, to balance contributions from clusters of different sizes. We then smoothed with a gaussian kernel (σ = 0.1 m), normalized the ISD to sum to 1, and subtracted the baseline ISD obtained from a uniform spike distribution in time. We computed the excess prominence, defined as the sum of all peak prominences between 0 m and 2.5 m in the ISD, adjusted for chance levels by subtracting the mean peak prominence sum obtained from shuffled data (*n* = 1,000 shuffles), where a random circular shift was applied to each flight trial between 0 s and the duration of the trial. For each cell we averaged the excess prominence at K over all combinations of K clusters, then took the median across cells and normalized to the median excess prominence at K=1 to obtain the normalized prominence score.

To determine if a plane-dependent grid code is sufficient to explain the observed reduction in periodicity when flights are pooled, we simulated grid cells (*n* = 360 cells) from hexagonal grid patterns defined on the plane of motion of each flight cluster, with consistent grid spacing, orientation, and phase relative to the reference frame spanned by its first two principal components with origin at the trajectory take-off position. For unclustered flights, the plane of motion was defined for each flight trajectory. For each simulated cell, grid spacing was uniformly sampled between 0.8 m and 1.5 m, orientation between 0° and 60°, and phase from the unit cell. Field sizes were defined as gaussian bumps with σ = 0.125 × spacing, and expected spike counts were randomly sampled from the spike count distribution of real cells. Simulated cells were redrawn if fewer than four fields landed within 2σ of a flight path in the pooled clusters. The same combinatorial flight cluster pooling procedure was performed on the simulated cells. As the unclustered flights already contain diverse flight trajectories, they were included only in the separate group containing all flights in a session.

Stability of grid cells (n = 236) was computed separately on clustered and unclustered flights using the Pearson correlation between 3D rate maps of odd / even flight trials. To account for differences in spike count and occupancy, we defined normalized stability for each cell as (*r*observed - *r*shuffle) / (*r*ceiling - *r*shuffle), where *r*observed is the observed rate map correlation between odd/even trial splits, *r*shuffle is that obtained from per-trial circular shifted spike shuffles, and *r*ceiling is estimated from sampling spikes directly from the combined rate map across all trials. The same analysis was applied to simulated cells with grid spacing and spike counts matched to the real grid cells, and σ = 0.125 × spacing. Non-random plane-referenced grid cells were simulated as described above with orientation and phase relative to the local planar reference frame. Random plane-referenced cells instead had random orientation and phase on each flight path, except for within a clustered flight path, where the orientation and phase were shared across trials. Room- referenced cells were simulated from HCP, FCC, and random field positions, with results pooled across the models. Shared segments were identified between pairs of clustered paths as the longest stretch along which their mean flight path remained within 0.3 m of each other, and heading direction remained ≤60° (same direction) or ≥120° (opposing direction). Similarity of spatial tuning on shared segments was assessed by normalized stability and across minus within correlation between the odd trials of one flight cluster with the even trials of same cluster (within- cluster) or other cluster (across-cluster), averaged across the different possible combinations. Shuffles were performed by circularly shifting all trials within one flight cluster (holding other cluster fixed) by the same random offset between 0.1 T and 0.9 T, where T is the trial duration; this was repeated for the other cluster and the results were averaged (n = 100 shuffles). Heading tuning was computed by binning spike counts and azimuthal heading direction (estimated by the tangent of the position trace smoothed at σ = 50 ms on samples with speed ≥ 1 m s^−1^) using 12° bins, and were circularly smoothed (gaussian kernel, σ = 1.5 bins). Heading direction tuning was assessed by the mean vector length (MVL) tested against the MVLs of simulated Poisson spike trains sampled from the cell’s 3D rate map (voxel size, 0.1 m; σ = 1.2 bins) which accounts for the expected heading direction tuning from the spatial firing pattern of the cell (n = 1000 shuffles, P < 0.05).

### Toroidal latent variable model

To rigorously assess whether the toroidal latent representations during clustered flights with clear grid cell periodicity generalize to that of unclustered flights despite the lack of periodicity, we adapted the toroidal latent variable model (LVM) from the architecture and code provided by Bjerke et al.^38^. In brief, the LVM consists of an encoder that maps population activity to a toroidal latent state, and a decoder that predicts neural activity from the latent state. The LVM is given by *p*(*x*, *z*) = *p*(*x*|*z*)*p*(*z*), where *x* is the population activity vector and *z* = (*z*_1_, *z*_2_) is the toroidal latent state. The decoder *p*(*x*|*z*) models the activity of cell i as *x_i_* ∼ *Poisson*(*g*(*z*, *u_i_*)), where *g*(*z*, *u_i_*) is a Gaussian-like tuning curve centered at the cell’s preferred phase *u_i_*, with Euclidean distances in sine/cosine coordinates on the toroidal manifold. The latent prior factorizes as *p*(*z*) = *p*(*z*_1_)*p*(*z*_2_) with (*z*_1_, *z*_2_) each drawn from a wrapped Normal distribution. The encoder is a 1D convolutional neural network (window size k = 5 time bins) parameterizing the variational posterior *q*(*z*|*x*), allowing the model to be trained end to end by maximizing a variational objective based on the evidence lower bound (ELBO). The analysis was performed on the population spiking activity, with no smoothing, of co-modular cells for each of n = 5 grid modules. All models were trained with the same parameters: ‘learning_rate’ = 2×10^−3^, ‘weight_kl’ = 0.1, ‘num_hidden’ = 256, ‘weight_time’ = 0, ‘batch_size’ = 256. To compare between clustered and unclustered flight conditions, we averaged across subsamples of each condition with the same duration (n = 20 subsamples). For each subsample we performed 5-fold cross-validation with one model trained on clustered flights only and another on unclustered flights only. For each fold, each LVM was trained on 80% of its respective subsample of flight time bins and evaluated on the same held-out 20% of unclustered flight time bins. For each cell, Pearson correlation was computed between the smoothed (σ = 30 ms) predicted and true spike counts. As a control, we applied a random circular shift to each cell’s spike trains (between 10% and 90% of its duration) on the clustered flights, and compared LVMs trained on shuffled clustered-flight data with LVMs trained on unshifted unclustered-flight data, both evaluated on the same held-out, unshifted unclustered time bins (*n* = 100 shuffles).

To evaluate whether the mapping of the latent structure onto spatial behavior learned by the LVM trained on clustered flights only matches that of an LVM trained on unclustered flights directly, we estimated toroidal phase maps from the latent toroidal coordinates of each model (see ‘Toroidal phase map estimation’). We then aligned the toroidal coordinates (see ‘Stability of toroidal tuning’) and computed the mean vector length (MVL) of the angular difference between their angular phase maps. To account for variations across different training seeds, each model fit and alignment was repeated 100 times, and the median was reported. Significance was evaluated with respect to a shuffle (*n* = 1,000), where the toroidal coordinates decoded by the LVM trained on clustered flights were circularly shifted by a random offset between 0 and the trial duration (same shift to both ***ϕ***1 and ***ϕ***2) on each unclustered flight trial, and the same toroidal alignment procedure was performed.

### Comparisons of alternative grid models

Columnar (COL), hexagonal close-packed (HCP), face-centered cubic (FCC), and 1D periodic time / distance models were fit by the log-likelihood of observed spikes under the respective model candidate rate maps grid searched across the range of parameters and fit separately for unclustered and clustered flights. For each cell, model fitting between unclustered and clustered flights was performed between each clustered flight path and unclustered flights, trial-matched by subsampling the flight trials of the larger group (average over n = 30 subsamples), z-scored against its shuffle, and averaged across flight clusters to yield a z-scored log-likelihood score. Shuffles were constructed by applying random per-trial circular shifts between (0, L), where L is the number of time bins in the flight trial (*n* = 100 shuffles). The COL model was fitted by searching over the 4D parameter space over spacing (30 steps log-spaced from 0.6 m to 2.5 m), orientation (15° steps), and 2D phase (8 × 8 = 64 steps), and modeled with cylindrical gaussian firing fields (σ = 0.125 × spacing). HCP and FCC lattices were searched spanning a 7D parameter space over spacing (30 steps log-spaced from 0.6 m to 2.5 m), 3D orientation (15° steps), and 3D phase (8 × 8 × 8 = 512 steps), and modeled with 3D gaussian firing fields (σ = 0.125 × spacing) arranged in HCP or FCC lattices. To account for symmetries, HCP lattice orientation range spanned 360° in azimuth, 90° in pitch, and 90° in roll, yielding ∼10 million total candidates across the 7D search space after deduplicating symmetric arrangements. FCC lattices spanned 180° in azimuth, 90° in pitch, and 90° in roll, yielding a total of ∼4.5 million candidates after deduplication. To evaluate the columnar model, we additionally evaluated the gridness of top-down activity on unclustered flights, which measures the six-fold symmetry expected of a columnar code. We computed the autocorrelation as done in ‘Alignment of plane-referenced grids’ using an annulus from 0.5 × s to 1.3 × s for each candidate spacing between 0.6 m and 2.5 m (75 steps log-spaced), and defined the columnar gridness as the maximum gridness obtained across this range to ensure that the low observed gridness was not due to failure in detection of autocorrelation peaks or spacing estimation. Significance was evaluated with respect to columnar gridness of shuffles generated by circularly shifting spikes on each flight trial between 0.1 T and 0.9 T, where T is the duration of each trial (n = 1000 shuffles).

1D periodic time and distance models were evaluated by linearizing firing rate along the trajectory in time (0.01 s bins) or in distance (0.05 m bins) and fit by a grid search over a 2D parameter space across spacing (75 steps log-spaced from 0.175 s to 0.729 s for time and 0.6 m to 2.5 m for distance) and phase (14 steps) and modeled by 1D gaussian firing fields arranged periodically in time or distance (σ = 0.125 × spacing). To account for phase offsets between flight trials, time-lagged (up to 0.5 s) or distance-lagged (up to 1.5 m) cross-correlation alignment was applied to align the activity on each trial to that of the longest flight trial with spikes. Trials shorter than 2 s or 6 m, and flight clusters with fewer than 5 trials, were excluded. To avoid alignment from reversing the circular shifts applied to shuffled data, a segment-shuffle procedure was performed where each trial was randomly segmented into three segments with minimum length of 0.5 s or 1.5 m and reordered non-cyclically (n = 100 shuffles). Note that while neural activity on clustered flights was already well-aligned without the alignment procedure, it received the same alignment procedure to ensure rigorous comparison between unclustered and clustered flights. To control for potential error accumulation during flight resulting in reduced periodicity on longer flights, we performed the same analysis using a length matched subset of unclustered flights to each clustered flight (within ±15% path length), and flight clusters with fewer than 5 length-matched unclustered flights were excluded.

### Detection of 3D lattice structure

To evaluate the presence of 3D lattice structure in the arrangement of firing fields when pooling together all flights, we followed similar procedures to those established in Ginosar et al.^18^ for comparability. In brief, rate maps were estimated by binning spikes and occupancy from all flights (both clustered and unclustered) into 15 cm cubic voxels. Binned spike and occupancy maps were adaptively smoothed by expanding each voxel to 1, 3 or 5 voxels, stopping whenever the expanded occupancy reached 300 ms, and voxels were omitted if occupancy did not reach the occupancy threshold. For peak detection only, rate maps were then smoothed with a gaussian kernel (σ = 1.5 voxels). Peaks greater than the mean rate map firing rate, and greater than the 75^th^ percentile of the value of the same voxel in shuffled rate maps (n = 2,000), were defined as firing fields. Firing fields within 0.6 m of each other were merged by keeping the one with higher firing rate. Shuffled rate maps were constructed by applying random circular shifts (between 10 s and T – 10 s, where T is the total duration of all flights) to the spike train. Stability of cells was defined by splitting data into odd and even flights in chronological order and computing the Pearson correlation between their respective 3D rate maps. A cell was included in the detection of 3D lattice structure if it (i) had an odd/even correlation of at least 0.3, (ii) had a minimum of 10 firing fields, and (iii) had at least 200 valid shuffles within ±3 firing fields of the observed field counts (n = 980 cells satisfied all three conditions).

Lattice fitting was performed by grid search over the 7D parameter space of hexagonal close-packed (HCP, ∼41 million candidates) and face-centered cubic (FCC, ∼6 million candidates) lattices, which are defined by spacing (0.60 m to 2.50 m in 40 geometrically spaced steps), 3D orientation (10° steps for HCP and 15° steps for FCC, deduplicated by symmetry), and 3D phase offset (8 steps per axis). Lattice fit quality was quantified by the root mean square (RMS) distance between observed field positions and their nearest lattice nodes. A cell was significantly fit by a lattice if its minimum RMS score was lower than the 5^th^ percentile of the shuffles (*n* = 200 valid shuffles, as described above).

### Lennard-Jones local order model

Modeling of the Lennard-Jones (L-J) local order model was performed following similar procedures as previously described in Ginosar et al.^18^. The L-J potential is defined by a characteristic interaction distance (r0) that controls the equilibrium distance between fields, and the temperature (T) that controls the thermal noise level which displaces fields away from their equilibrium position (Fig. S16A). The local order model posits that the inter-field-distance (IFD) of grid cell firing fields in volumetric 3D space is governed by pairwise interactions described by the L-J potential. To simulate field positions under the L-J model, N fields were initialized at random positions within the visited occupancy of the room and iteratively relaxed towards lower energy configurations using Metropolis Monte-Carlo simulations where a candidate displacement to each field position was iteratively proposed. Candidate displacements were accepted if the resulting field energy Ei is lower than previous field energy, and otherwise with probability exp(- ΔE/T). As configurations require more steps to converge for lower T, the number of relaxation steps was chosen based on T (10^5^ steps at T = 0; 5×10^4^ steps at T ≤ 0.25; 2×10^4^ steps at T < 1; 10^4^ steps at T < 10; 2×10^3^ steps at T ≥ 10). Simulations of the L-J model were evaluated and compared to observed data based on the inter-spike-distance distribution (ISD), which serves as a proxy for inter-field distance when firing fields are well defined without the need to detect firing fields. The ISD is computed by binning all spike-pair distances with 0.05 m bins from 0 to 5 m, then smoothed with a gaussian kernel with σ = 0.03 m. ISD of simulated field arrangements was estimated from the spike positions sampled from observed occupancy under the simulated fields, with gain set to match the expected simulated spike counts to observed data. Simulated fields were modeled as 3D gaussians with σ = 0.125 × d, where d is the median nearest-neighbor distance of the resulting field arrangement. Simulated ISDs were evaluated by the normalized fraction of achievable similarity score, S, defined as

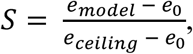

where *e_model_* = *RMS*(*ISD_model_* − *ISD_observed_*), *e_ceiling_* = *RMS*(*ISD_ceiling_* − *ISD_observed_*), and *e*_0_ = *RMS*(*ISD_observed_*). The achievable error *eceiling* is estimated by the model computed from the rate map estimated directly from observed spike data without smoothing, sampled multinomially to achieve exact spike counts (median of n = 300 draws). RMS errors are compared against the reference of predicting no spikes, given by *e0*.

The free parameters of the L-J model were fit via a grid search over r0 (39 steps from 0.50 m to 2.40 m), and T (24 steps including T = 0, quarter-decade steps from 10^−2^ to 10^1^, and half-decade steps from 10^1.5^ to 10^6^), with 1,000 arrangements simulated at each grid step. For each cell, the (r0, T) that yielded the minimal median RMS error to the observed ISD (evaluated from 0.6m to 2.5 m) on the combined activity over all flights, both clustered and unclustered, was chosen. The fitted model was then evaluated on each clustered flight path separately using the median similarity score S across 1,000 simulations (scored over the 0.6 m – 4 m ISD band). As the ISD is sensitive to the number of spikes sampled on each flight cluster, we included only flight clusters with a minimum of 75 spikes, and excluded cells that did not have any flight cluster with sufficient spikes (keeping n = 255 out of 331 grid cell candidates). For comparison, we fit the plane-dependent grid model (2D-Hex) with a similar random-sampling based procedure. First, the grid spacing was fit by a grid search (49 steps from 0.60 m to 2.52 m) by sampling 1,000 arrangements (each with random phase and orientation) on the best-fit plane for each clustered flight path, selecting the spacing that yielded the lowest average of the median RMS across flight clusters. The fitted model was then evaluated on clustered flights using the same normalized similarity score and 0.6 m to 4 m ISD band as the L-J model over the 1,000 random arrangements.

### Data Analysis

All analyses were conducted using custom code in Python and MATLAB (R2024b, MathWorks). The following open-source packages were used: numpy (1.26.4), scipy (1.16.1), scikit-learn (1.9.0), scikit-image (0.26.0), ripser (0.6.12), rastermap (1.0), pytorch (2.11.0), kilosort4 (4.1.1). Computationally heavy workloads were performed on the Savio computational cluster provided by the Berkeley Research Computing program at the University of California, Berkeley.

### Statistical analysis

Sample sizes were not predetermined using formal methods; sample sizes used were comparable to those of similar studies. Experiment sessions were not randomized, and experimental conditions were not blinded. All statistical tests were two-sided unless otherwise stated. Empirical permutation tests, which compare observed statistics with a shuffle distribution, and binomial tests were one-sided.

## Acknowledgments

We thank A. Vollan for guidance on grid cell recordings with Neuropixels, and B. Dunn, E. Hermansen, and V. Olsen for discussions on analysis techniques for grid cell population activity. We also thank I. Nelken and members of his lab (M. Karayanni & M. Jankowski) for initial guidance on procedures for Neuropixels implants. We thank the Yovel lab for making the dataset of Egert-Berg et al. (2021) publicly available.

## Author contributions

K.K.Q. and M.M.Y. designed the research. K.K.Q. performed experiments. K.K.Q. and M.M.Y. analyzed the data and wrote the manuscript.

---

**Figure S1.**
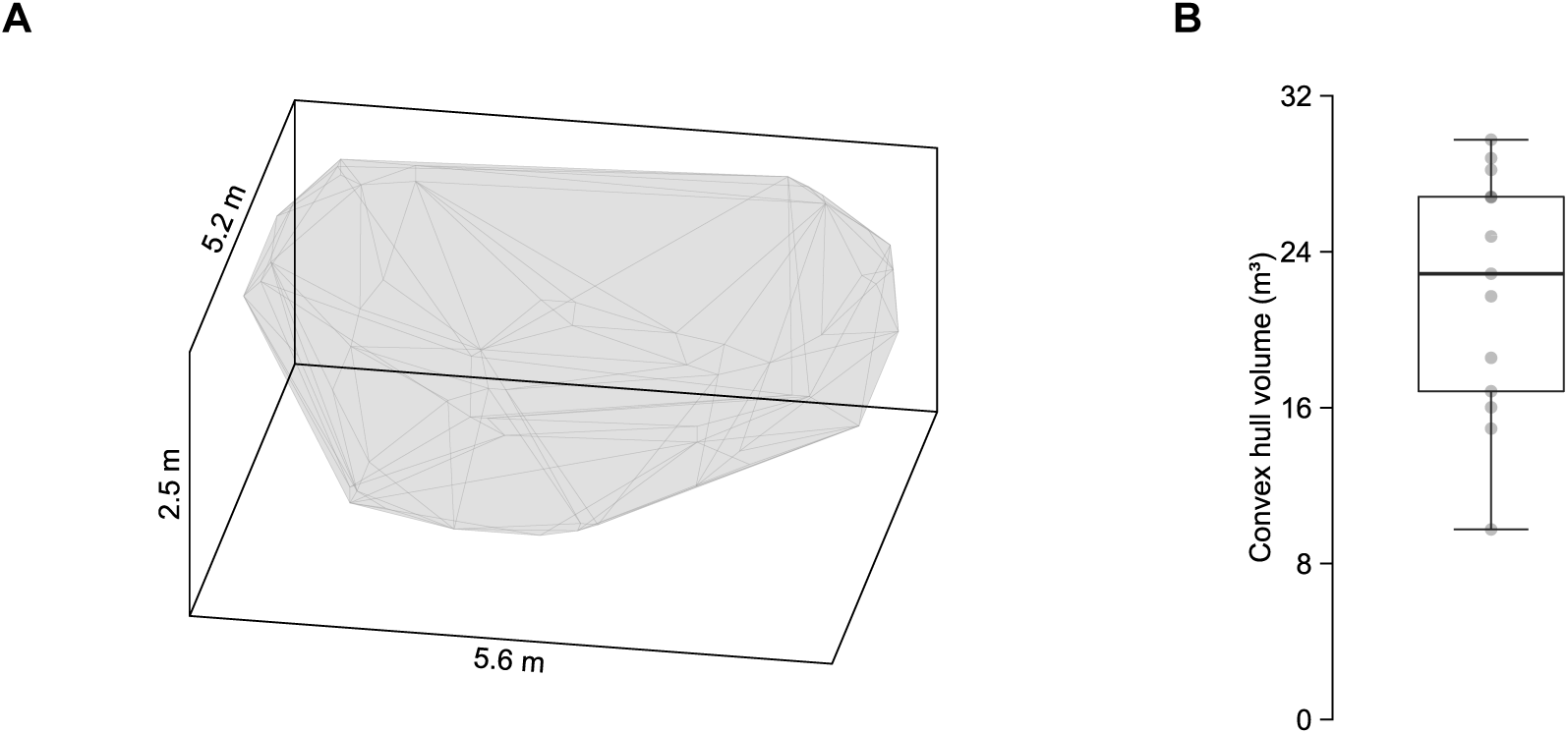
Three-dimensional volumetric flight behavior. **(A)** Convex hull of flight trajectories during an example session, constructed by binning position into 15 cm voxels and taking the convex hull of occupied-voxel centers. (**B**) Volume of convex hull of flight trajectories across sessions (*n* = 13). Box plot shows the median (center line), 25^th^ to 75^th^ percentiles (box bounds), and minimum and maximum values (whiskers). Volume for each individual session is indicated with gray circles.

**Figure S2.**
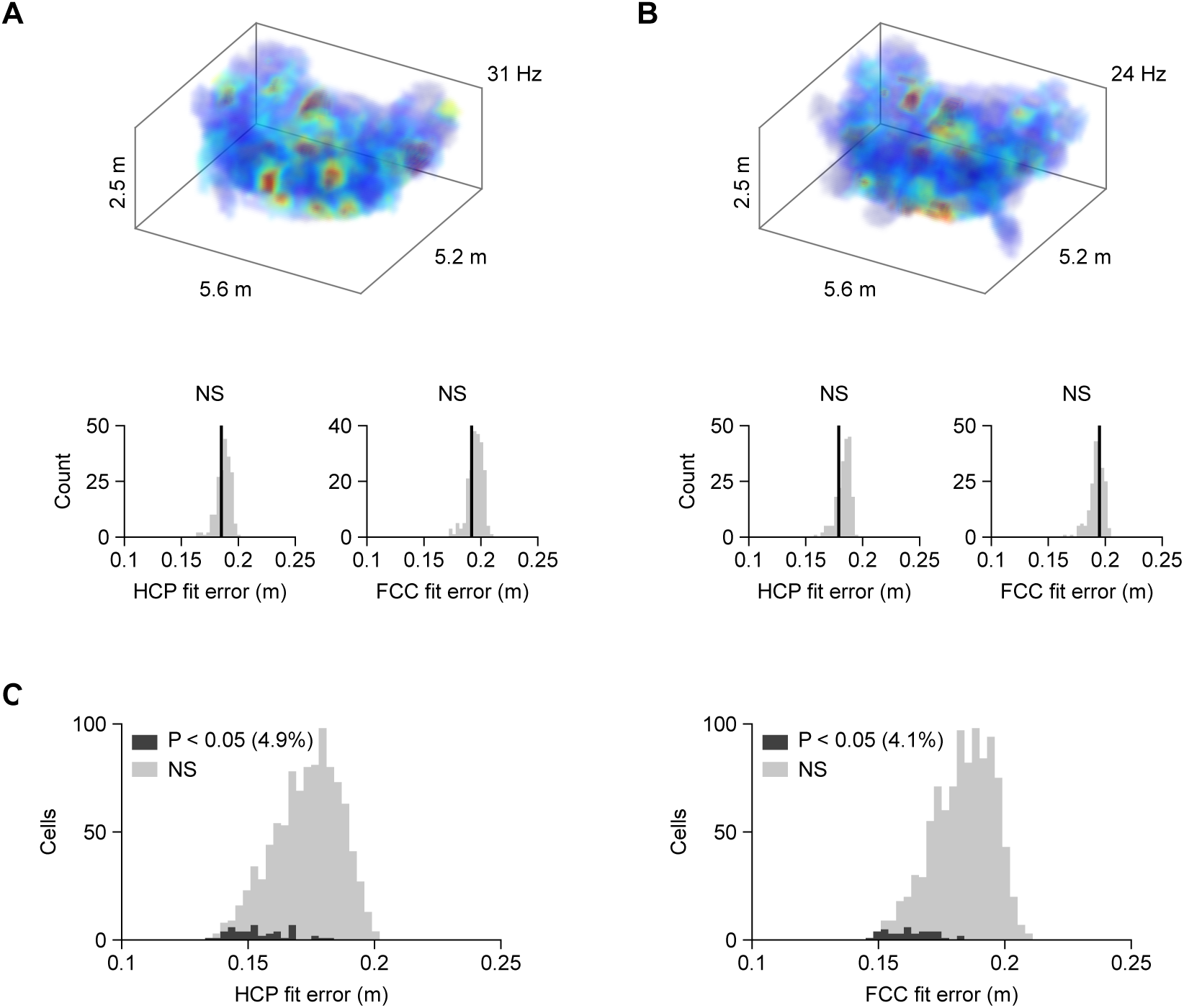
Lack of global 3D lattice structure when pooling together all flights. Quantification of lattice structure during flight when pooling together all flights, following similar procedures as established in Ginosar et al.^18^ (Methods). (**A** and **B)** Quantification of lattice structure for two example cells when all flights are pooled together. Top, 3D rate map, with the color scale maximum (indicated) set to the 99^th^ percentile of firing rates. Bottom, lattice fit under hexagonal close-packed (HCP, left) and face-centered cubic (FCC, right) 3D lattices for observed activity (black line) versus distribution of shuffled fits (gray). NS, not significant. (**C**) Histogram of HCP (left) and FCC (right) lattice fits for the subset of cells with 10 or more firing fields when pooling all flights in a session, and stable between odd and even flights (*n* = 980). Dark gray shading indicates cells with significant lattice fits (chance level 5%), light gray shading indicates non-significant lattice fits (NS).

**Figure S3.**
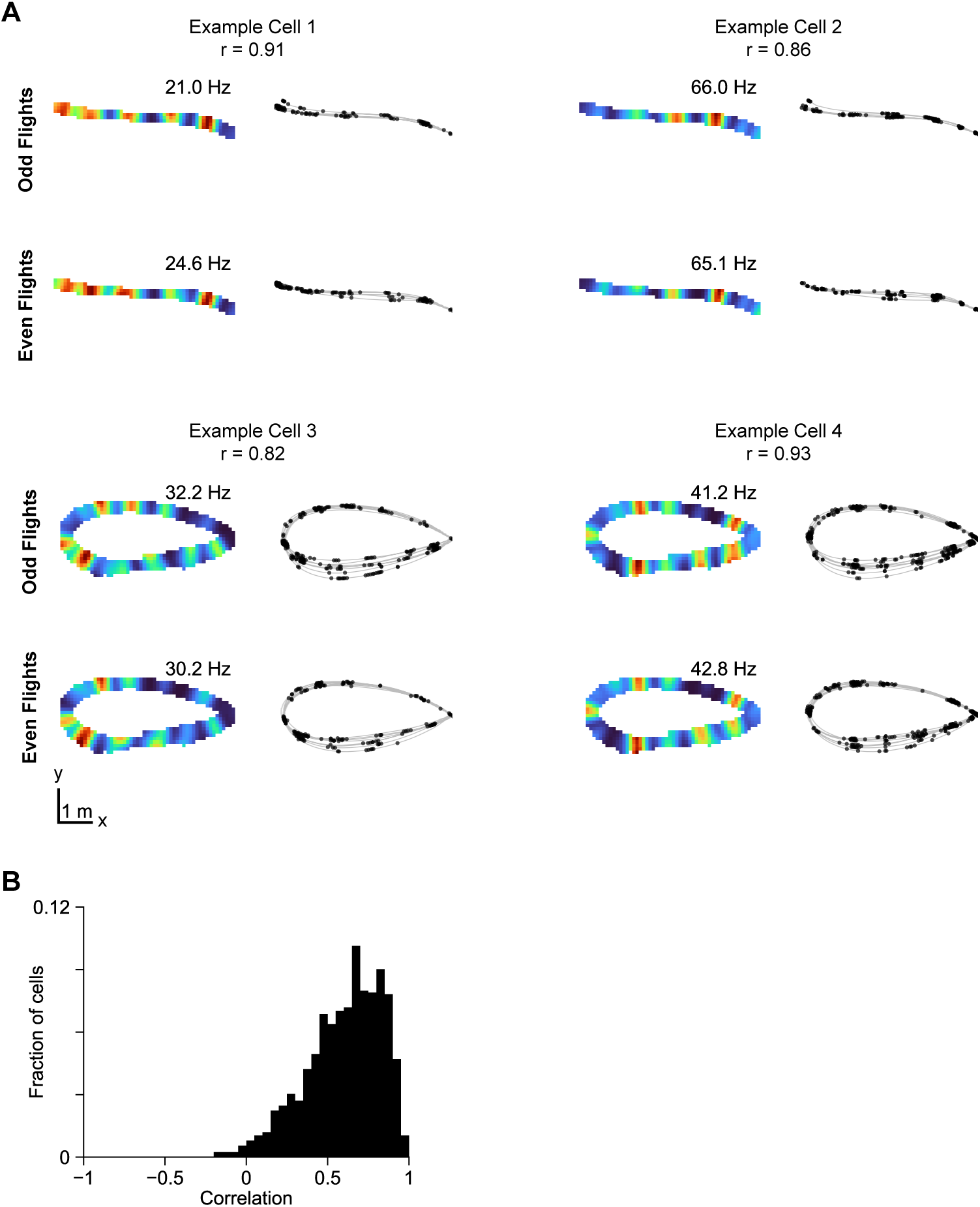
MEC neurons exhibit stable spatial firing fields across flights. **(A)** Rate maps and raw spike locations for four example MEC neurons along repeated flight trajectories, split by odd and even flights. For each example, occupancy-normalized firing-rate maps are shown on the left and spike locations (black) overlaid on flight trajectories (gray) are shown on the right. Peak firing rate for each rate map is indicated; r, the odd-even rate map Pearson correlation. (**B**) Distribution of correlation between odd- and even-flight rate maps across cells meeting the spike count and field count criteria (≥ 75 spikes and ≥4 fields on at least one clustered flight; n = 1,253).

**Figure S4.**
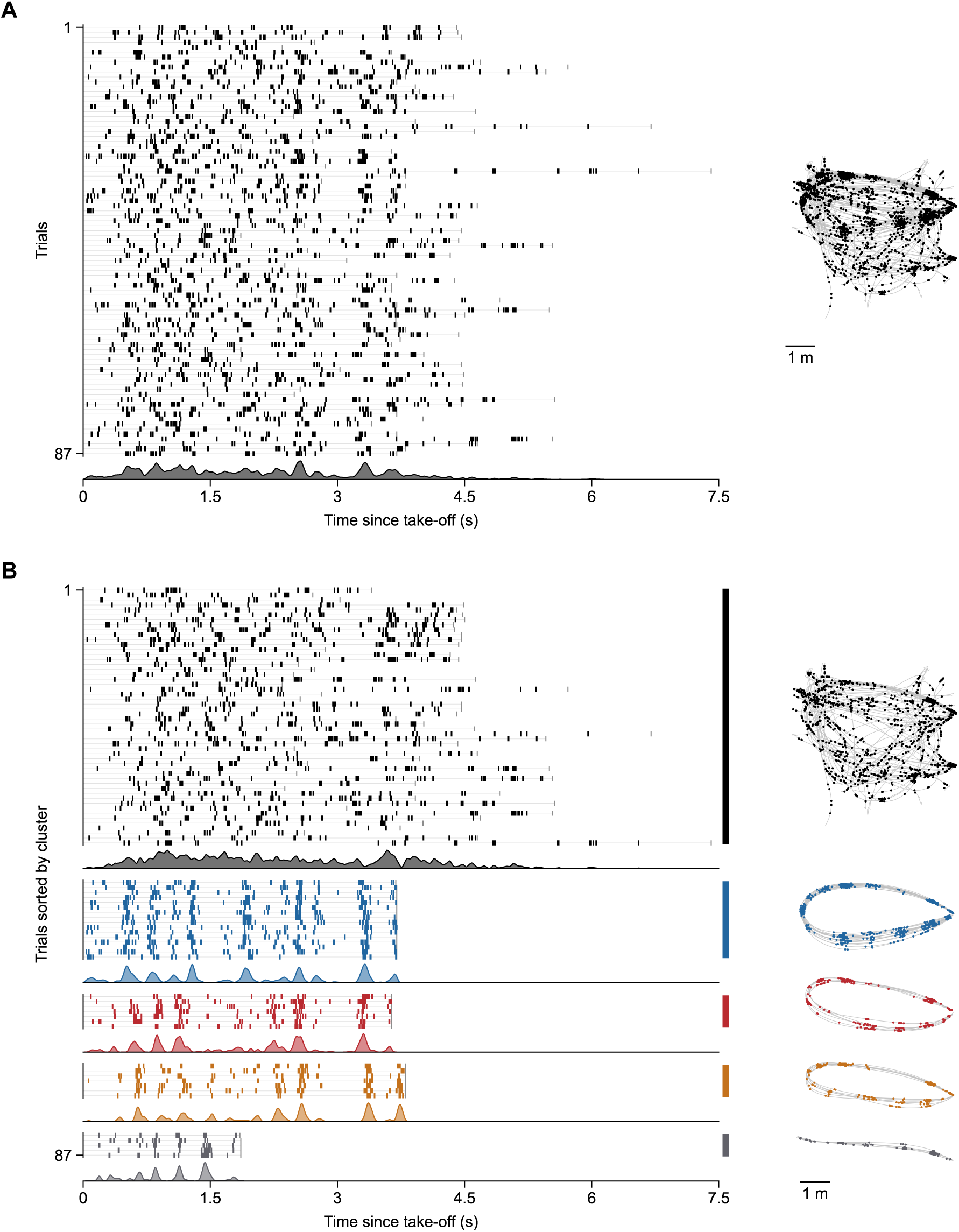
Sorting spike raster by flight cluster reveals periodic firing patterns. (**A**) Left, spike raster plot aligned to flight take-off with random trial ordering of an example cell during all flights. Horizontal gray lines and gray tick indicate duration of flight for each trial. Right, corresponding spikes (black) plotted on position (gray). Bottom, peristimulus time histogram (PSTH) of the spikes in the corresponding raster, peak-normalized for visualization. (**B**) Same example cell and behavior as in **A**, but spike raster and PSTH are sorted and colored by flight cluster (left). Right, corresponding spikes (colored by flight cluster matching that of the spike raster) on position (gray) separated by flight cluster.

**Figure S5.**
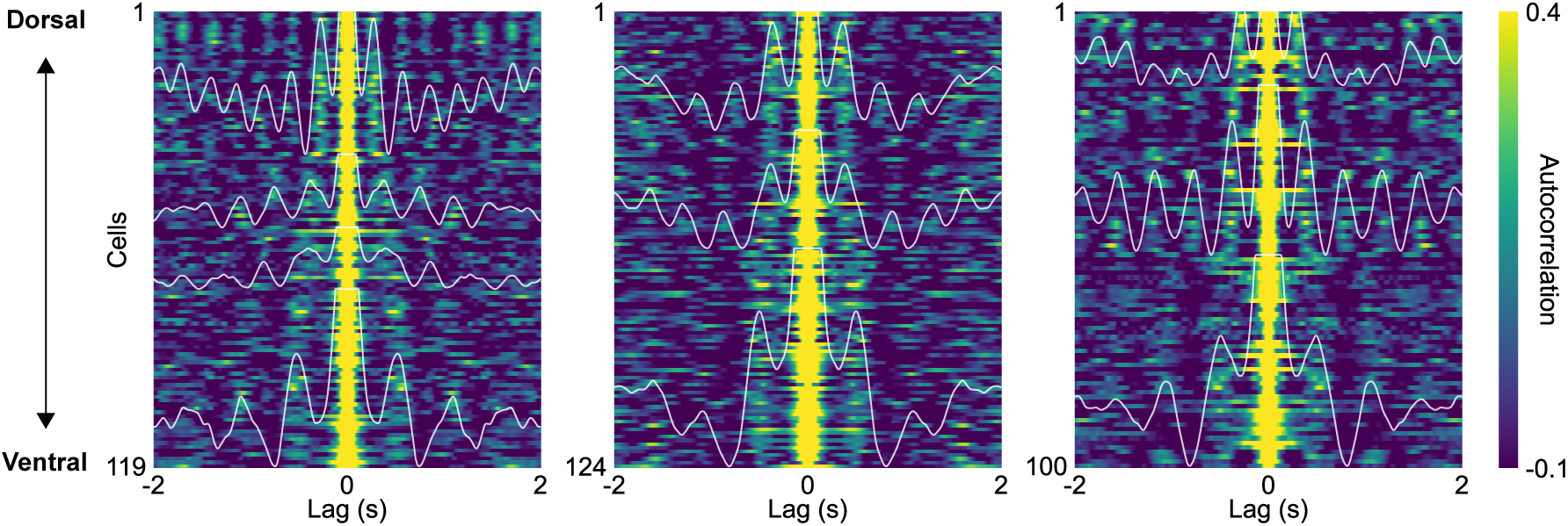
Topographic module-like organization of grid cells along dorso-ventral MEC axis. Time-lag autocorrelations of significantly periodic grid cells, shown separately for each bat and sorted from dorsal-to-ventral by mean cluster recording depth. Cells were clustered using the normalized power spectral densities (PSD) of their firing rates across all bats. White traces indicate the mean autocorrelation for each cluster. Only within-bat clusters containing 15 or more cells are shown.

**Figure S6.**
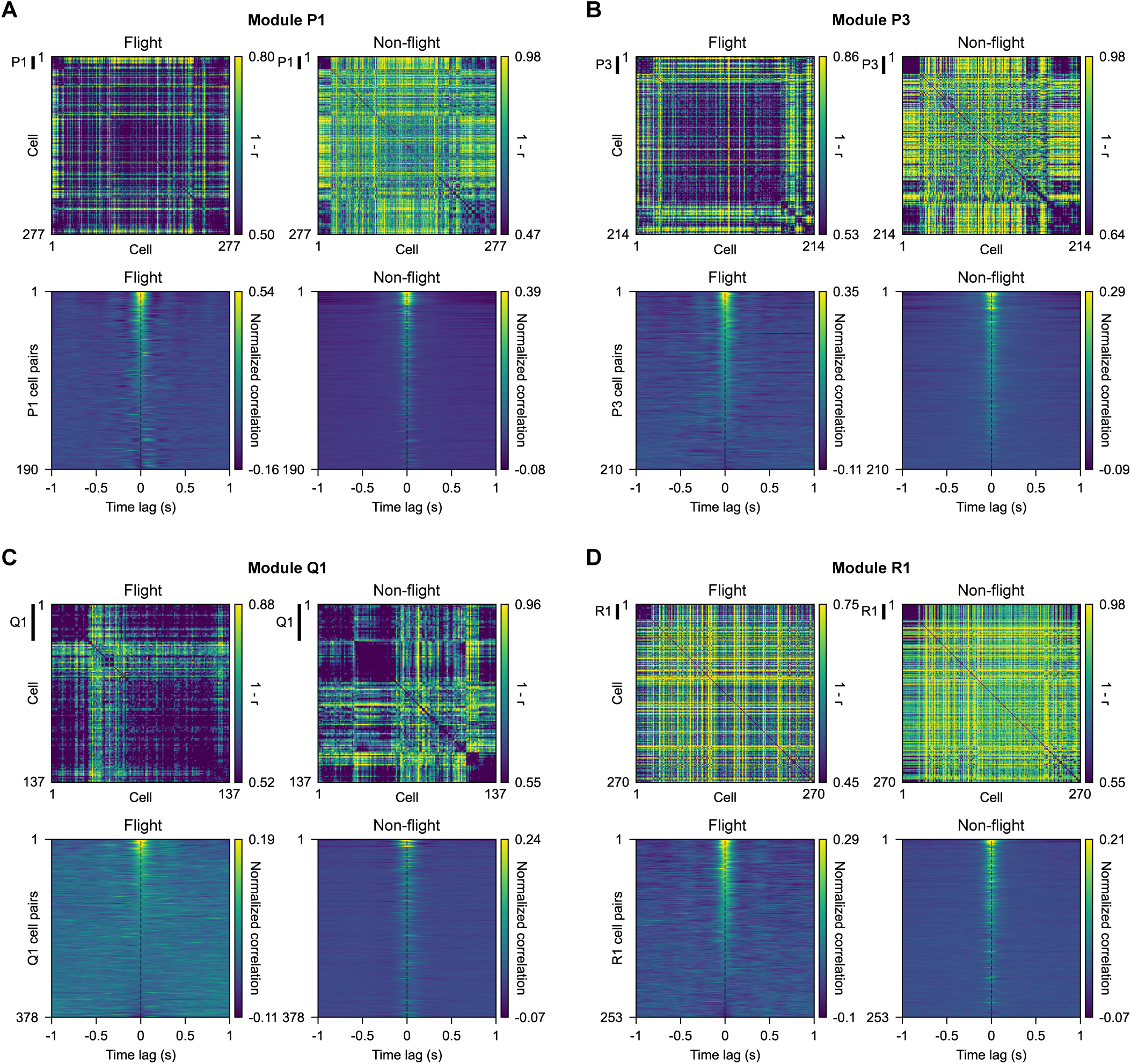
Stable pairwise spike-time correlations of co-modular grid cells across behavioral states. (**A-D**) Spike-time correlation-based clustering of four additional putative grid modules not shown in Fig. 2A,B, computed separately during flight and non-flight bouts. For each module, top panels show correlation (*r*) distance matrices during flight (left) and non-flight (right) periods, sorted identically according to grid module clustering. Bottom panels show within-module pairwise normalized cross-correlation during flight (left) and non-flight (right) periods, sorted by zero-lag correlation during flight bouts. The same sorting was applied to non-flight bouts. The cross-correlation of each cell pair was normalized by mean subtraction.

**Figure S7.**
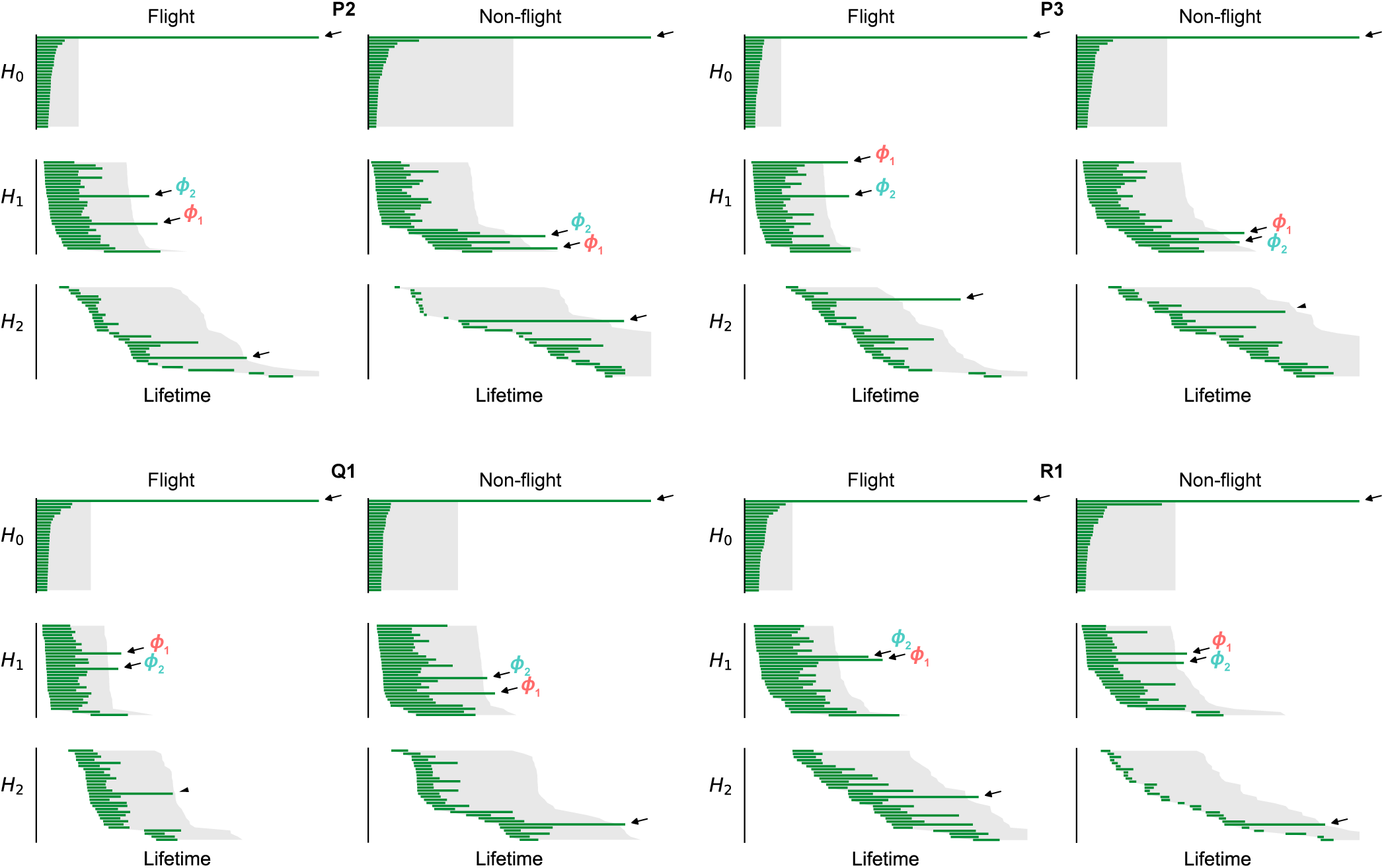
Persistence barcodes indicating toroidal topology of grid cell population activity. Persistence barcodes computed during flight (left) and non-flight bouts (right) for the four additional putative grid modules not shown in Fig. 2C: P2, P3, Q1 and R1. Green bars indicate the lifetime of topological features in each homology dimension: connected components (H。), circular features (H,) and cavities (H,). Arrows mark the prominent features, which are consistent with a toroidal topology. Gray shading indicates the 95^th^ percentile of the shuffle distribution, constructed by circularly shifting each cell’s spike train relative to the others. Non-significant, but prominent feature lifetimes are indicated instead with a triangle (P3 non-flight H_2_, P = 0.10, lifetime = 1.6× 2nd longest H_2_ bar; Q1 flight H_2_, P = 0.066, lifetime = 2.0× 2nd longest H_2_ bar). The test statistic was the longest bar for H。 and H,, and the second-longest bar for H,.

**Figure S8.**
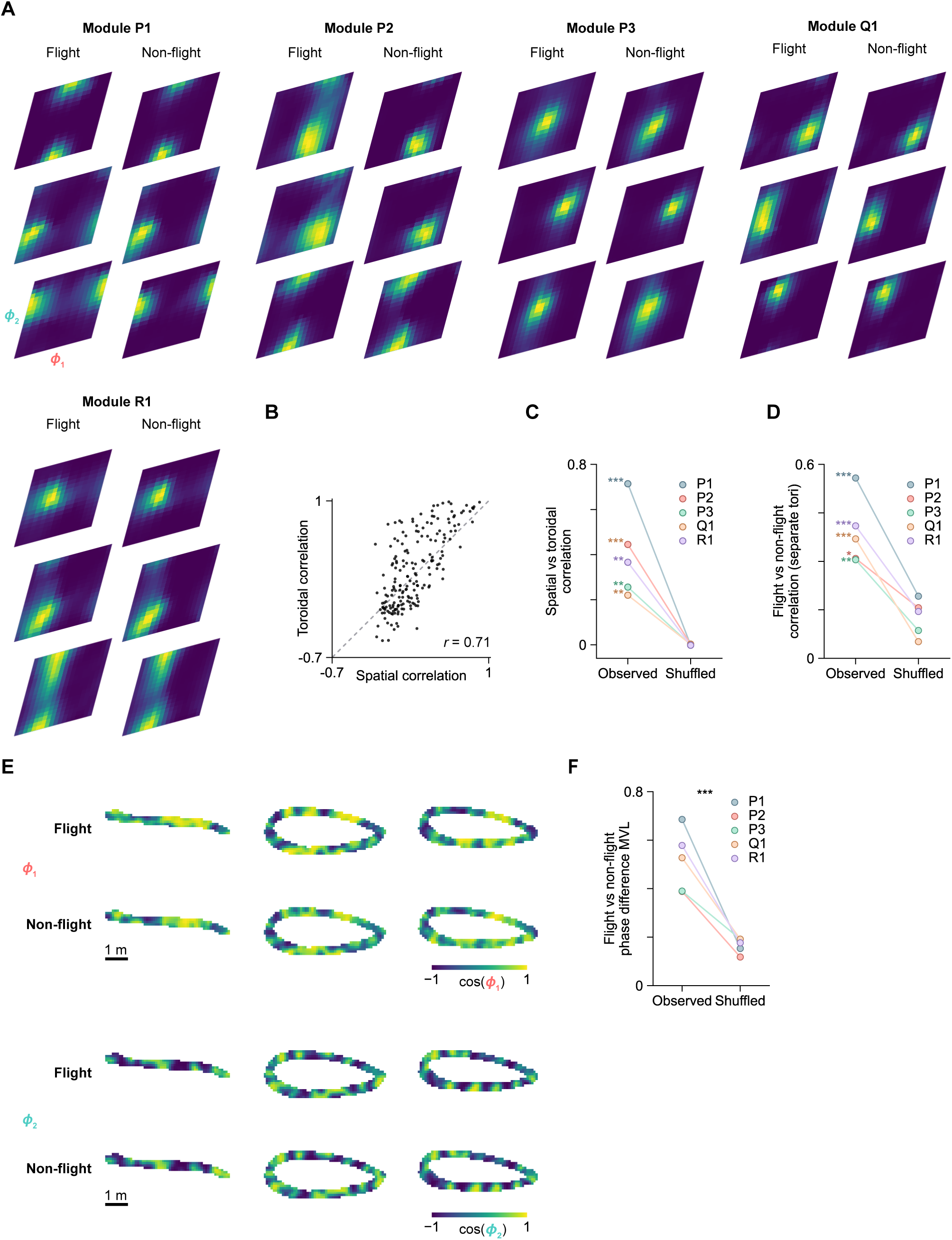
Toroidal rate maps are stable across flight and non-flight bouts. **(A)** Example toroidal rate maps of grid cells from each module during flight (left), and non-flight bouts (right). Toroidal rate maps are plotted in toroidal coordinates and normalized to the peak firing rate of each cell (Methods). (**B**) Scatter plot of pairwise toroidal rate map correlations versus pairwise spatial rate map correlations of the P1 module, constructed from independent time bins with non-flight bouts for toroidal rate maps and flight bouts for spatial rate maps (n = 190 cell pairs, Pearson r = 0.71). Gray dashed line, unity line. (**C**) Pearson correlation between pairwise spatial correlation and toroidal correlation, as shown in **B**, for all five grid modules compared to shuffle distribution constructed by random permutation of unit identity. n = 190 (P1), 190 (P2), 210 (P3), 378 (Q1), 253 (R1) cell pairs. ***P < 0.001, **P < 0.01. (**D**) Pearson correlation between flight and aligned non-flight toroidal rate maps, computed using separate toroidal parameterizations for each behavioral state (Methods). Lines connect paired comparisons for each module, indicated by colors. *n* = 20 (P1), 20 (P2), 21 (P3), 28 (Q1), 23 (R1) cells. \*\*\**P* < 0.001; P2: \**P* = 0.017, P3: \*\**P* = 0.002. (**E**) Example toroidal phase maps of P1 module of *ϕ*_1_ (top two rows), and *ϕ*_2_ (bottom two rows) decoded from separate toroidal parameterizations during flight (first and third rows) and non-flight (second and fourth rows). Non-flight torus is detected using only non-flight time bins, and toroidal coordinates are decoded on held-out flight bout time bins. Scale bar 1 m. (**F**) Mean vector length (MVL) of angular differences between toroidal phase maps, on flight clusters with at least 5 flights, decoded from separate toroidal parameterizations during flight and non-flight bouts for each module. Shuffles are constructed by applying the same random circular shift to *ϕ*_1_ and *ϕ*_2_ on each flight trial (decoded from non-flight torus), between 0 s and the trial duration, and recomputing the shuffled toroidal phase maps (*n* = 1,000 shuffles). For each shuffle, the same toroidal alignment is performed as done for observed data. \*\*\**P* < 0.001 for all modules.

**Figure S9.**
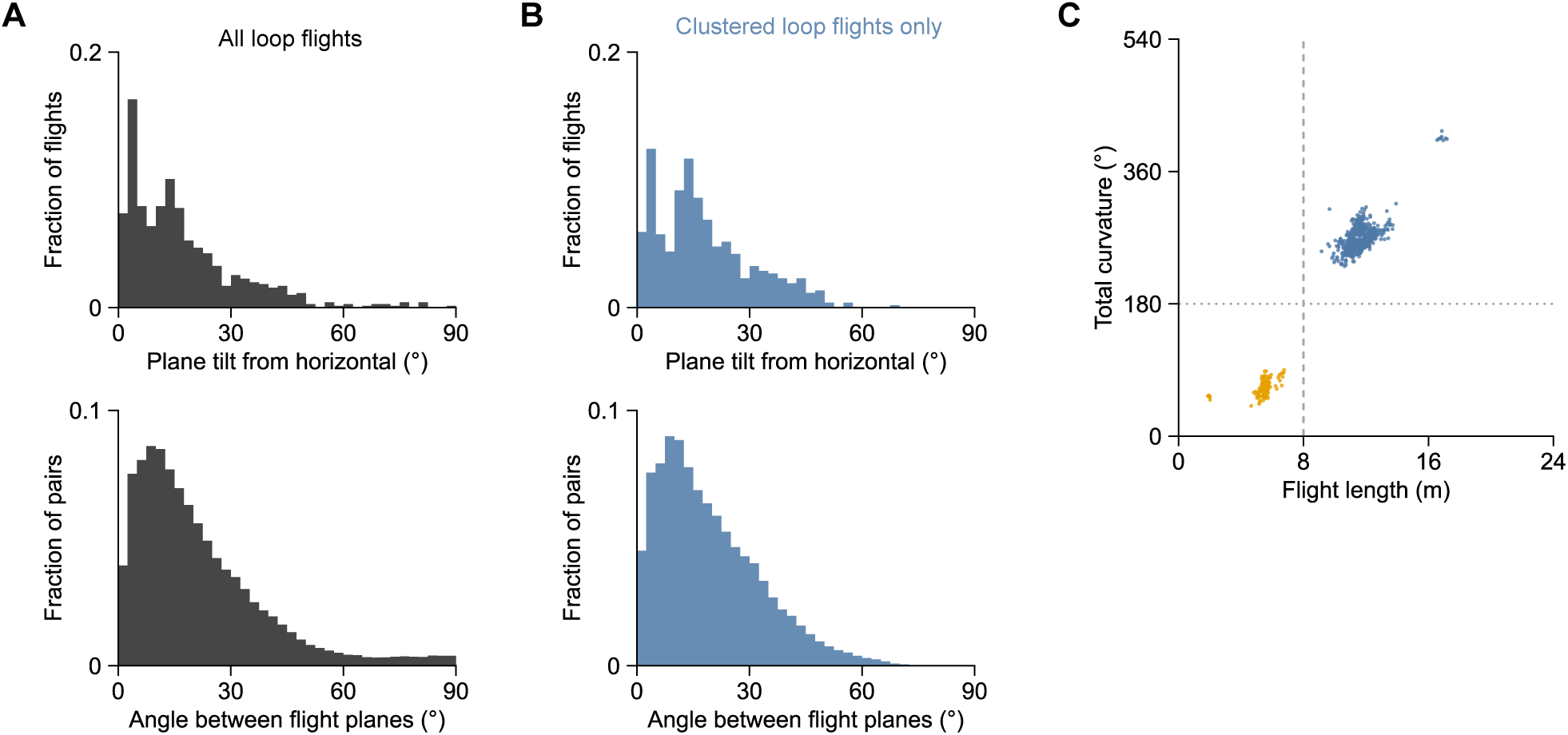
Orientations of laboratory flight planes. **(A)** Distribution of flight plane angles from horizontal (top) and between planes (bottom) for all unclustered and clustered laboratory non-straight loop flights (*n* = 705 flights). Plane angle is defined as the angle between the plane normal and the z-axis. (**B**) Distribution of angles from horizontal (top) and between planes (bottom), as in **A**, for clustered loop flights only (*n* = 523 flights). (**C**) Scatter plot of total flight curvature versus length of all clustered flights (n = 662 flights) showing a bimodal distribution corresponding to longer loop flights (blue) and shorter straight flights (orange). Gray dashed line, 8 m cut-off reliably separates loop and straight clustered flights. Gray dotted line, 180° total curvature, corresponding to reversal of flight direction. Total curvature was estimated by summing the turning angles of successive chords of the flight trajectory, resampled at 50 cm arc lengths and excluding the final chord corresponding to landing. When selecting for clustered loop flights (n = 523) which have longer length (median length = 11.49 m, IQR 11.1 m to 12.0 m) and higher total curvature (median curvature = 268°, IQR 258° to 278°), we found that 97.7% (511 of 523 flights) were well fit by a single plane that explains >98% variance, consistent with the results of Fig. 3E. Unclustered loop flights (between 180° to 360° curvature and 8 m to 16 m length) were similarly well described by a single plane (116 of 122 loop flights, 95.1%). Flights towards feeders (median tilt 14.5°, IQR 5.8° to 25.4°) varied more widely than flights towards self-selected perch locations (median tilt 8.9°, IQR 4.6° to 14.8°).

**Figure S10.**
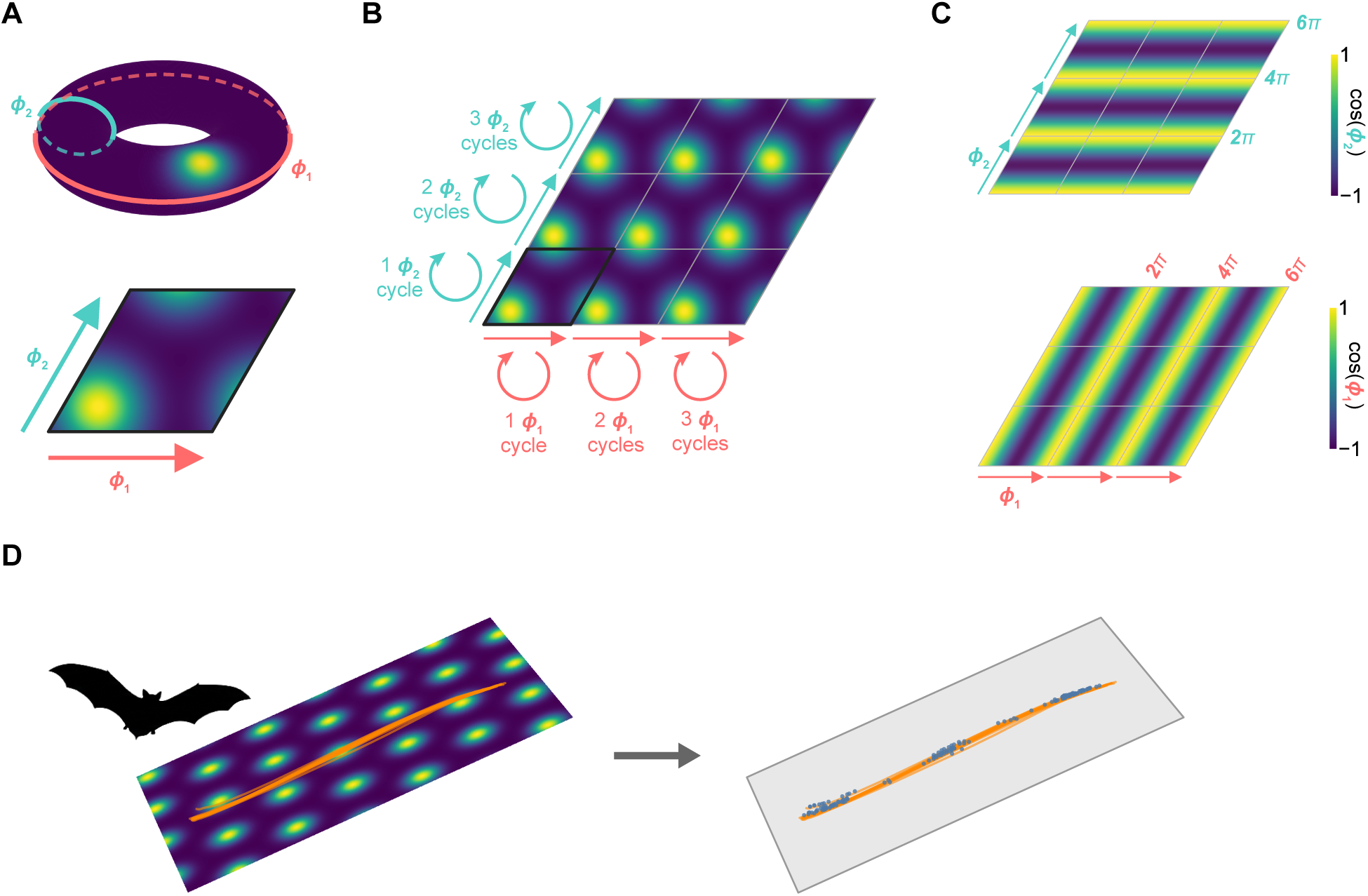
Conceptual illustration of mapping toroidal grid-cell representations onto two-dimensional space. (**A**) Top, schematic of a grid cell toroidal manifold defined by two circular coordinates, *ϕ*_1_ (orange) and *ϕ*_2_ (teal). Example tuning field denotes the activity of one grid cell at a specific *ϕ*_1_ and *ϕ*_2_ position on the torus. Bottom, the primitive rhombic unit cell of a two-dimensional hexagonal lattice. The torus is equivalent to this unit cell under periodic boundary conditions, with *ϕ*_1_ and *ϕ*_2_ spanning the two edges of the rhombus. (**B**) Periodic tiling of two-dimensional space by the rhombus unit cells generates a hexagonal grid pattern. *ϕ*_1_ and *ϕ*_2_ repeat along the principal axes of the hexagonal lattice, along the edges of the rhombi. (**C**) Toroidal phase coordinates (*ϕ*_1_, *ϕ*_2_) form plane-wave patterns across space, corresponding to the tiling of the rhombus unit cell. (**D**) Schematic of a bat’s flight trajectory (orange) intersecting hexagonal grid fields on its plane of motion (left). Under this mapping, firing (blue dots) along the flight trajectory is predicted to be consistent with the underlying hexagonal lattice (right).

**Figure S11.**
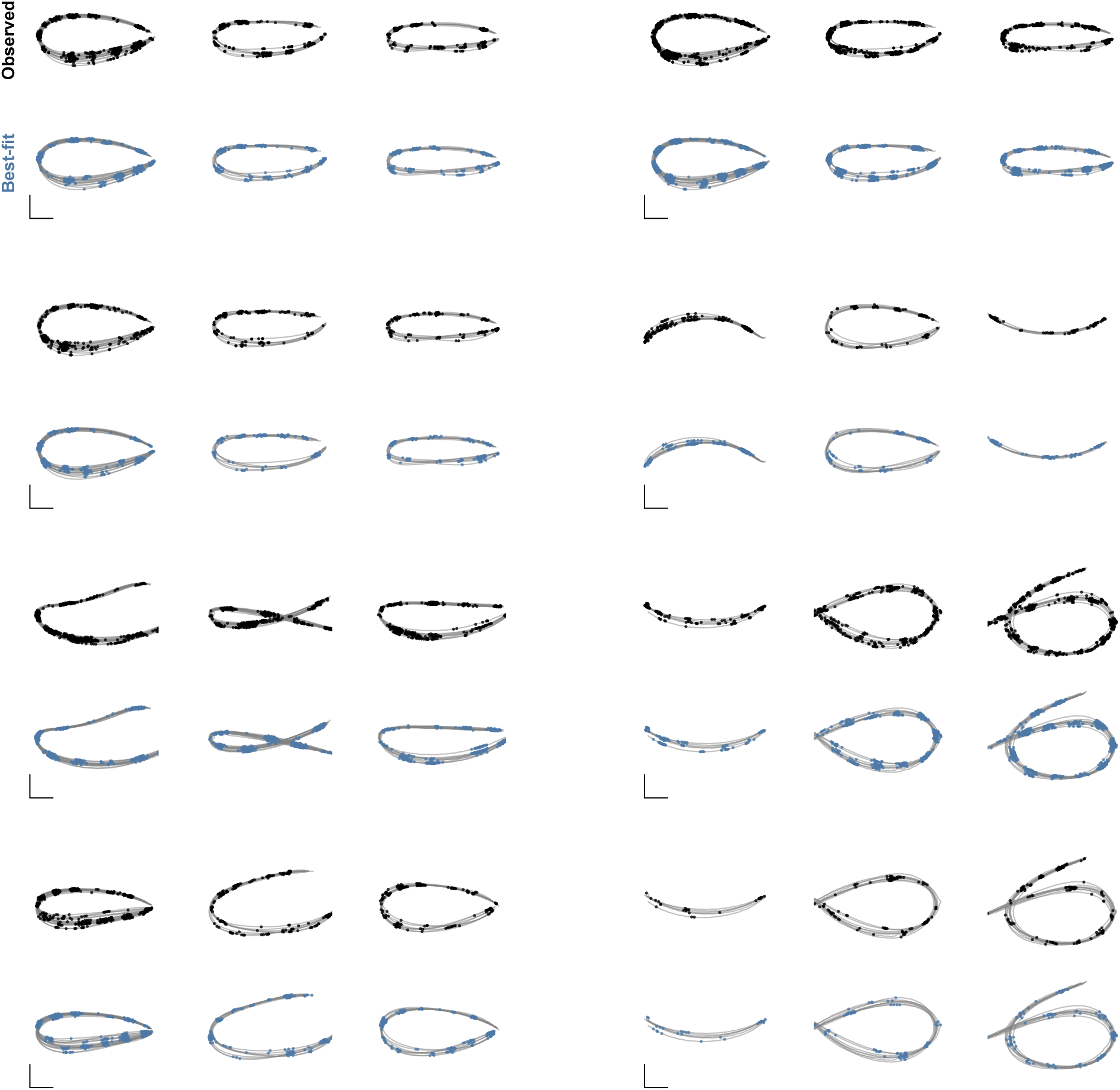
Additional examples of grid fitting on the plane of motion. Hexagonal grid fitting results for eight example grid cells on three clustered flight paths each. Black dots indicate observed spike positions overlaid on flight trajectories (gray). Blue dots indicate simulated spike positions from the best-fit two-dimensional hexagonal grid defined on the plane of motion. Scale bar, 1 m, in plane of motion coordinates.

**Figure S12.**
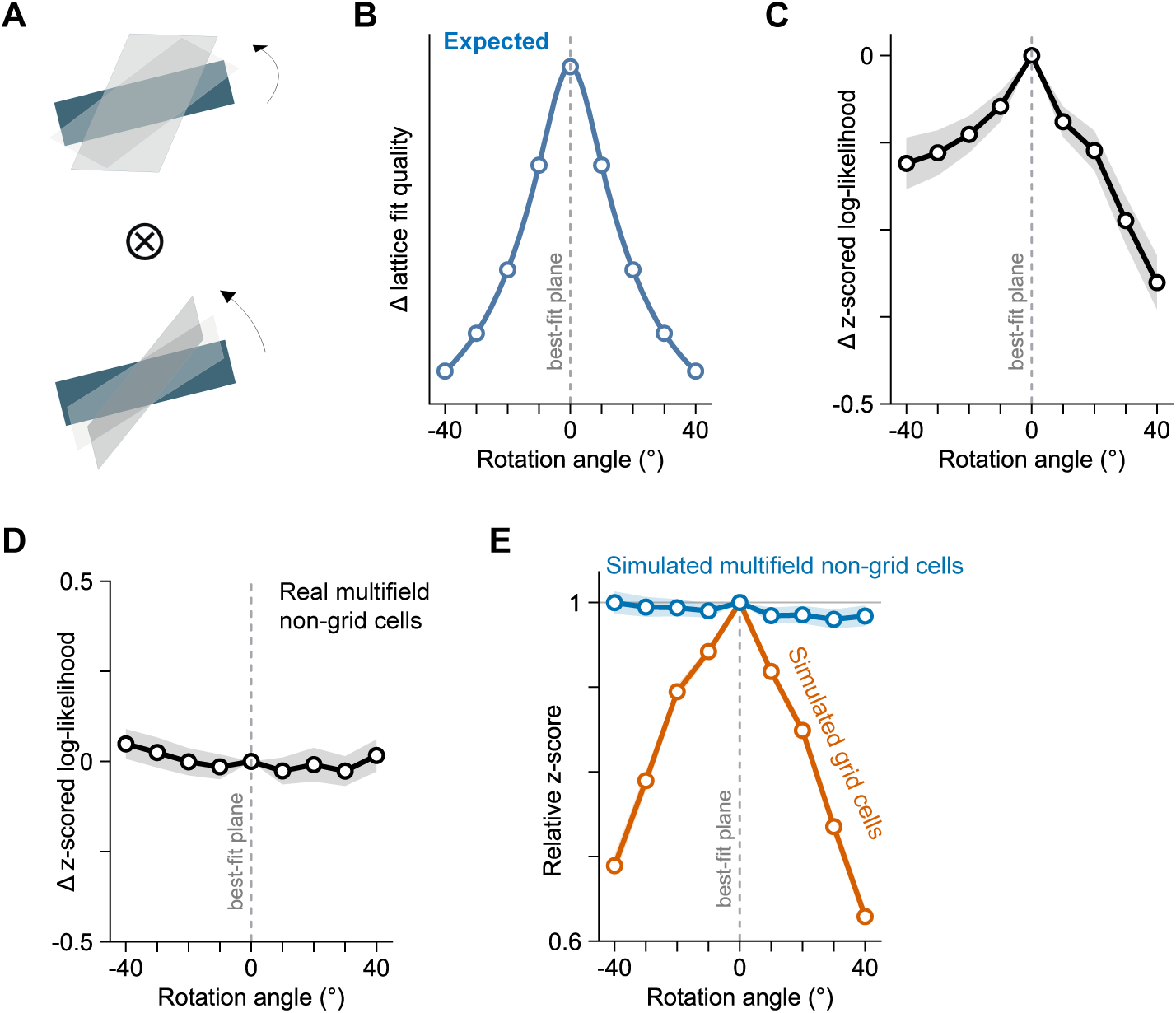
Grid fit quality decreases on planes rotated away from the plane of motion. **(A)** Schematic of candidate planes generated by rotating the best-fit plane of motion in azimuth and pitch, as shown in Fig. 3L. (**B**) Expected decrease in lattice fit quality on rotated planes if grid-cell representations are specifically aligned to the plane of motion, as shown in Fig. 3M. (**C**) Observed change in z-scored log-likelihood across rotated planes, as in Fig. 3N, for grid cells with significant Bonferroni-corrected grid fits on at least one of the 49 planes (average slope of lattice-fit reduction *β* = -0.062 ± 0.010 (mean ± s.e.m.) *z*-score / 10°, Wilcoxon signed-rank, *W* = 10016, *n* = 273 cells, *P* = 2.91 × 10^−11^). This inclusion criterion avoids selection bias from defining significant grid cells only on the best-fit plane. (**D**) Change in *z*-scored log-likelihood across rotated planes for non-grid multifield cells (average slope of lattice-fit reduction *β* = 0.003 ± 0.012 *z*-score / 10°, Wilcoxon signed-rank, *W* = 842, *n* = 58 cells, *P* = 0.92), defined as multifield cells without significant lattice fits on any plane. (**E**) Relative z-score, normalized by *z-*score on the best-fit plane, on rotated planes for simulated grid cells and non-grid multifield cells with matched field counts. Simulated grid cells exhibited significantly steeper reduction in lattice fit compared to simulated non-grid cells (average difference in slope Δ*β* = -0.077 ± 0.002 relative *z*-score / 10°, Wilcoxon signed-rank, *W* = 322, *n* = 360 pairs, *P* = 1.37 × 10^−59^).

**Figure S13.**
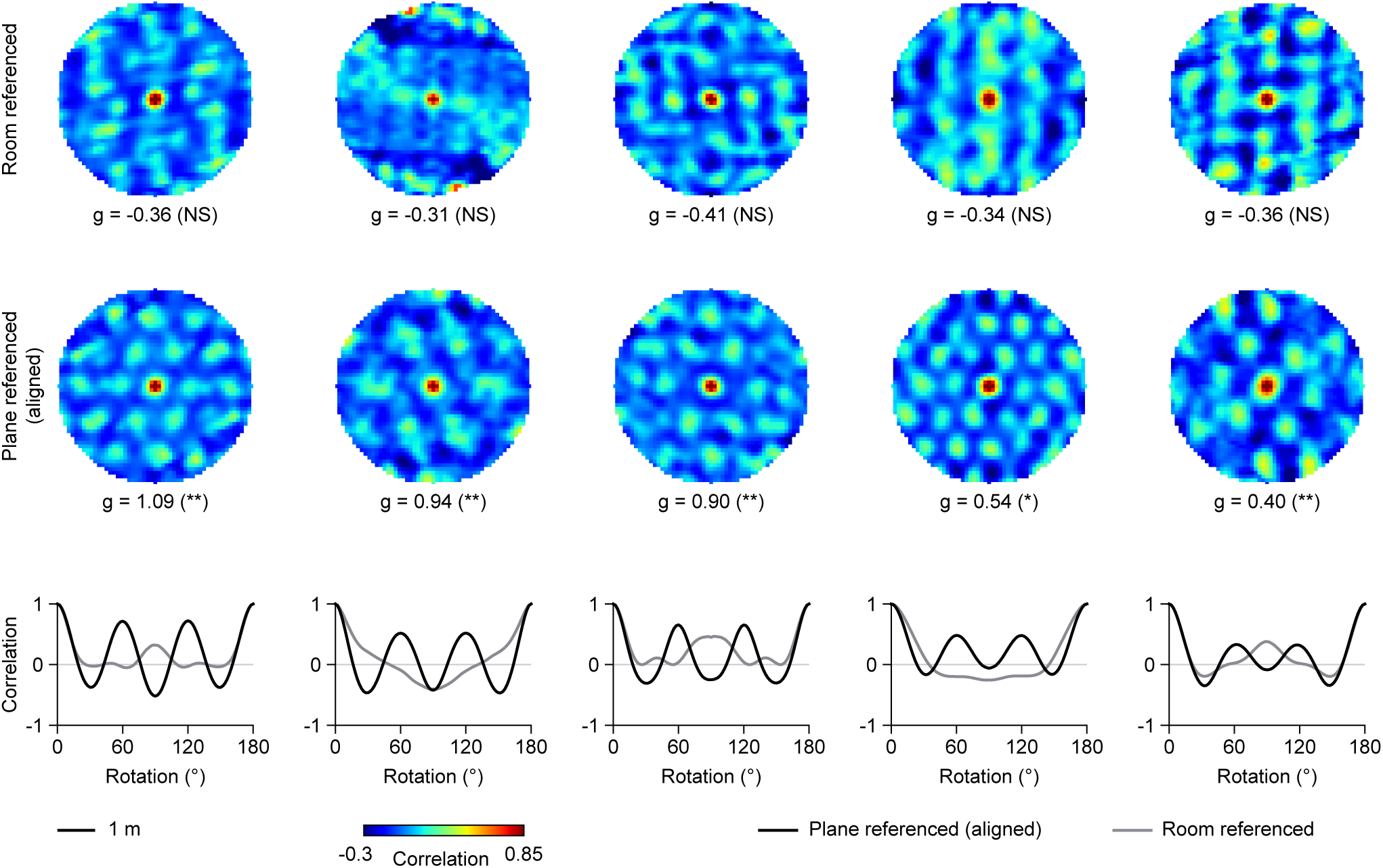
Alignment in plane-referenced coordinates reveals hexagonal grid patterns. Spatial rate-map autocorrelations of five example cells (columns) not shown in Fig. 4A computed in the room-referenced top-down projection (top row), and aligned plane-referenced shared coordinates (middle row). Cells shown are, from left to right, ranks 6, 9, 12, 37, and 41 among the 46 significant cells in aligned gridness. Gridness score, g, is indicated below each spatial autocorrelation with significance in parentheses (NS, not significant; \*\**P* < 0.01; \**P* < 0.05). Bottom, rotational autocorrelation computed on a circular annulus of the aligned plane-referenced (black) and room-referenced (gray) autocorrelation map. The annulus is centered at the central peak with inner and outer radii of 0.5× and 1.3× the estimated grid spacing, respectively. Significance is computed for each cell relative to its own shuffles (*n* = 1,000) in each reference frame, constructed by applying a random circular shift to each flight trial. Each shuffle underwent the same spacing estimation and alignment procedure before comparing with aligned gridness scores. A significantly higher number of cells had significant gridness in aligned plane-referenced coordinates (46 of 136, or 33.8%) compared to in room-referenced coordinates (9 of 136, or 6.6%; McNemar exact *P* = 3.02 × 10^−9^). All spatial autocorrelation maps share the same scale bar (1 m) and color scale (-0.3 to 0.85).

**Figure S14.**
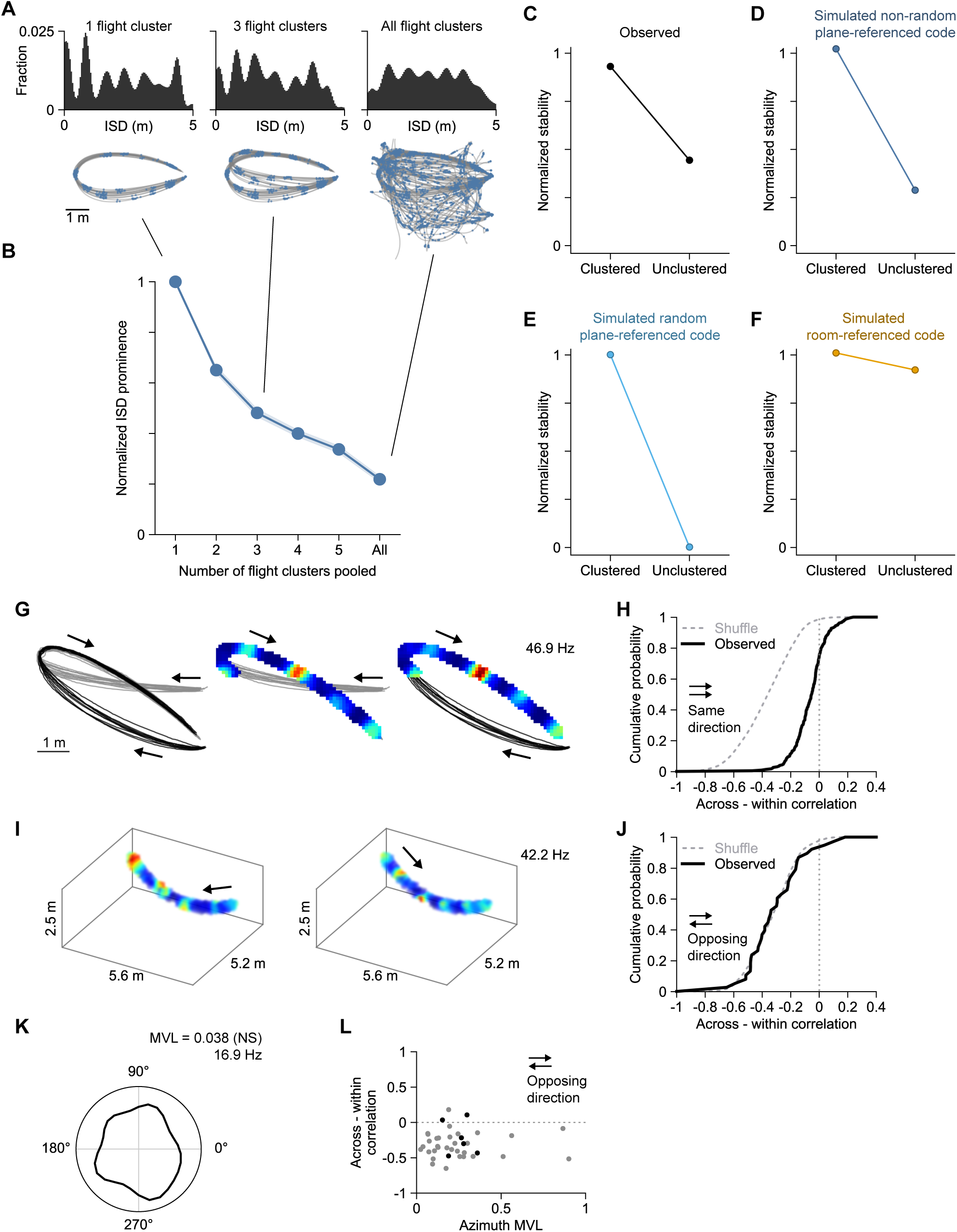
Grid cell activity across mixtures of flight paths. (**A**) Example simulated plane-referenced grid-cell response for one (left), three (middle), or all (right) flight clusters. Simulated grid cells were defined on the plane of motion of each flight trajectory (Methods). Top, inter-spike-distance (ISD) distribution. Bottom, simulated grid cell spikes (blue) over flight trajectories (gray). (**B**) Median ISD normalized prominence score of simulated plane-referenced grid cells (blue trace) as increasing numbers of flight clusters are combined. Shaded region indicates the bootstrapped standard error of the median. (**C-F**) Median normalized stability on clustered flights (left) versus unclustered flights (right). To account for differences in spike count and occupancy, we defined normalized stability for each cell as (*r_observed_* - *r_shuffle_*) / (*r_ceiling_* - *r_shuffle_*), where *r_observed_* is the observed rate map correlation between odd/even trial splits, *r_shuffle_* is that obtained from per-trial circular shifted spike shuffles, and *r_ceiling_* is estimated from sampling spikes directly from the combined rate map across all trials. (**C**) Observed grid cells (n = 236 cells with significant grid fits). (**D**) Non-random plane-referenced grid model where the spacing is fixed, grid phase is set relative to take-off, and grid orientation is set relative to PC1×PC2 local reference frame of the flight path. (**E**) Random plane-referenced grid model, where grid phase and orientation are random on every flight plane. (**F**) Room-referenced grid code in a fixed spatial frame. (**G**) Example pair of flight clusters (left) that shares a flight segment. Rate map of an example cell on the shared flight segment between the flight clusters (middle and right). Rate map voxels are shown only for the overlapping voxels between the pair, trajectories are overlaid for reference. Arrows indicate flight direction. Peak firing rate indicated. (**H**) Cumulative distribution of the across-cluster minus within-cluster correlation difference on shared segments traversed in the same direction (n = 234 cells, three bats; normalized stability, 0.93 ± 0.03, median ± s.e.). Similarity of spatial tuning was assessed by correlating the odd trials of one flight cluster with the even trials of same cluster (within-cluster) or other cluster (across-cluster). Black, observed. Gray dashed line, shuffles computed by circularly shifting each trial of one cluster by the same offset, between 0.1×T and 0.9×T, where T is the duration of the flight. (**I**) Example cell activity on a pair of flight clusters traversing the same shared segment in opposing directions. Arrows indicate flight direction. (**J**) Cumulative distribution as shown in **H** on shared segments traversed in opposing directions for a subset of the cells in **H** (n = 38 cells, one bat; normalized stability, 0.05 ± 0.07, median ± s.e.). (**K**) Azimuthal heading direction tuning curve of the example cell shown in **I**, computed over all unclustered flights. NS, not significant. Peak firing rates are indicated. (**L**) Scatter plot of across minus within correlation during opposing traversals versus mean vector length during unclustered flights of the same cells in **J**. Spearman ρ = 0.001, P = 0.99, n = 38 cells. NS, not significant. Black circles, significant MVL. Light gray circles, not significant MVL.

**Figure S15.**
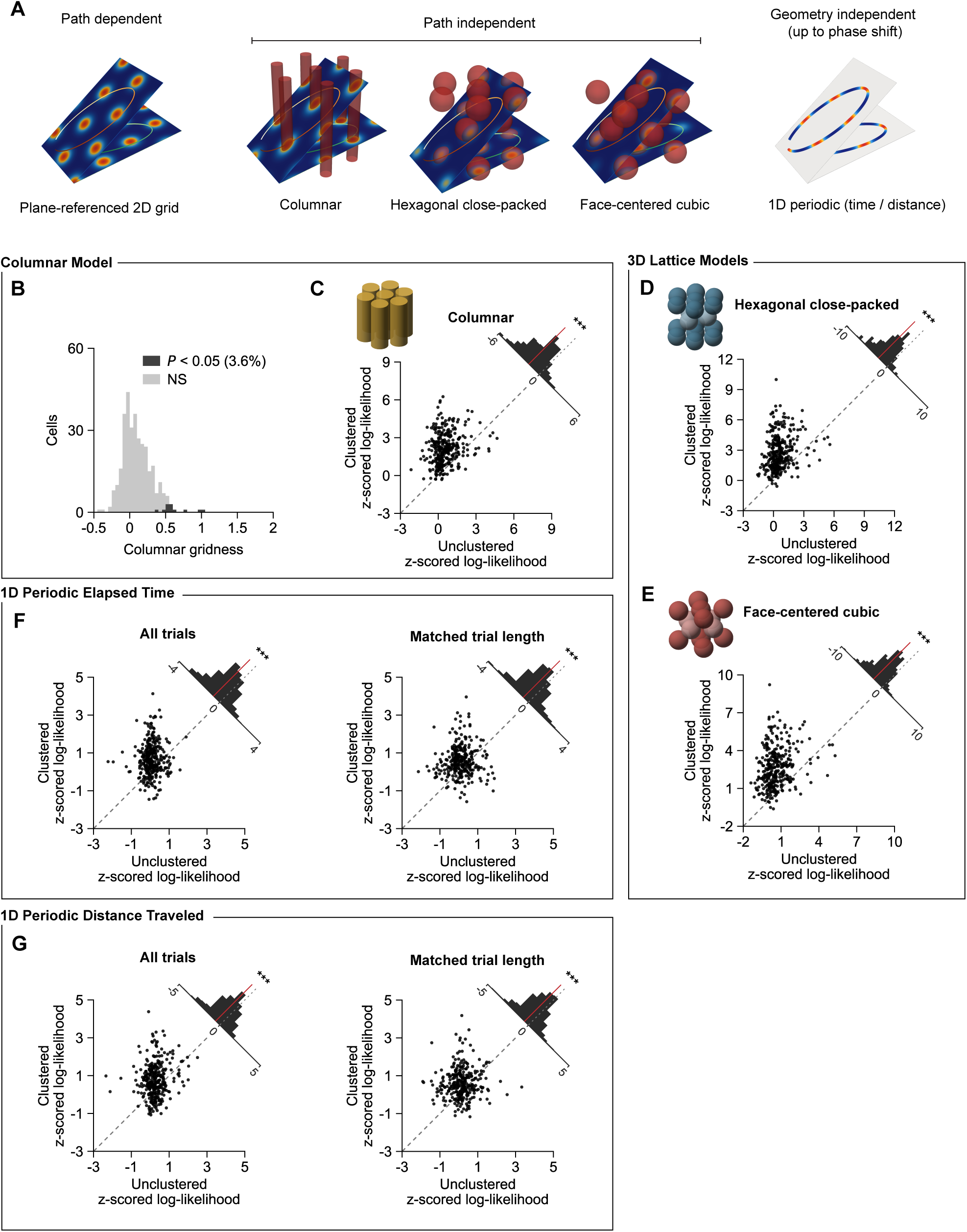
Alternative non-plane-referenced grid models. We assessed the possibility that periodic activity on clustered flight paths could arise from alternative non-plane-referenced grid models. (**A**) Schematics of the plane-referenced 2D grid model and the non-plane-referenced grid models. (**B**) Distribution of columnar gridness, computed by projecting activity during unclustered flights onto the horizontal plane. Dark shaded region, significant (12 of 331 cells, 3.6%; Binomial P = 0.90). Gray shaded region, not significant (NS). Under columnar (COL), hexagonal close-packed (HCP), and face-centered cubic (FCC) grid models, the activity of grid cells is a function of the bat’s position and is independent of the flight path the bat takes, which predicts comparable periodicity on both clustered and unclustered flights. 1D periodic distance or time codes predict consistent periodic activity along flight paths, up to a phase shift, irrespective of flight path geometry. (**C**-**G**) Paired scatter plot of z-scored log-likelihoods under each grid model on clustered flights versus unclustered flights. Dashed gray line, unity line. Inset, histogram of unclustered minus clustered z-scored log-likelihood differences. Red line indicates difference distribution mean. *N* = 331 cells, two-sided Wilcoxon signed-rank test. (**C**-**E**) In contrast to the predictions of the path-independent lattice models, consistency of single-cell activity with COL, HCP, and FCC lattices was significantly lower on unclustered flights than on clustered flights. (**C**) Columnar model. \*\*\**P* = 4.65 × 10^−46^, *W* = 2649. (**D**) Hexagonal close-packed model. \*\*\**P* = 9.78 × 10^−50^, *W* = 1639. (**E**) Face-centered cubic model. \*\*\**P* = 4.53 × 10^−50^, *W* = 1549. (**F**-**G**) Likewise, periodicity as a function of elapsed time or distance traveled was significantly lower on unclustered flights than on clustered flights, despite 1D periodic models being independent of flight path geometry. Trial-to-trial neural activity was aligned based on time- or distance-lagged cross-correlation to account for potential phase shifts. Comparison was performed using all trials as well as trials of matched lengths to account for potential confounds due to accumulation of error during flight. (**F**) 1D periodic time code, using all trials (left, \*\*\**P* = 9.33 × 10^−25^, *W* = 9574), and subset of trials with matched trial lengths (right, \*\*\**P* = 4.60 × 10^−17^, *W* = 10301, *n* = 304 cells). (**G**) 1D periodic distance code, using all trials (left, \*\*\**P* = 3.46 × 10^−17^, *W* = 12785), and matched trials (right, \*\*\**P* = 1.92 × 10^−14^, *W* = 11437, *n =* 304 cells).

**Figure S16.**
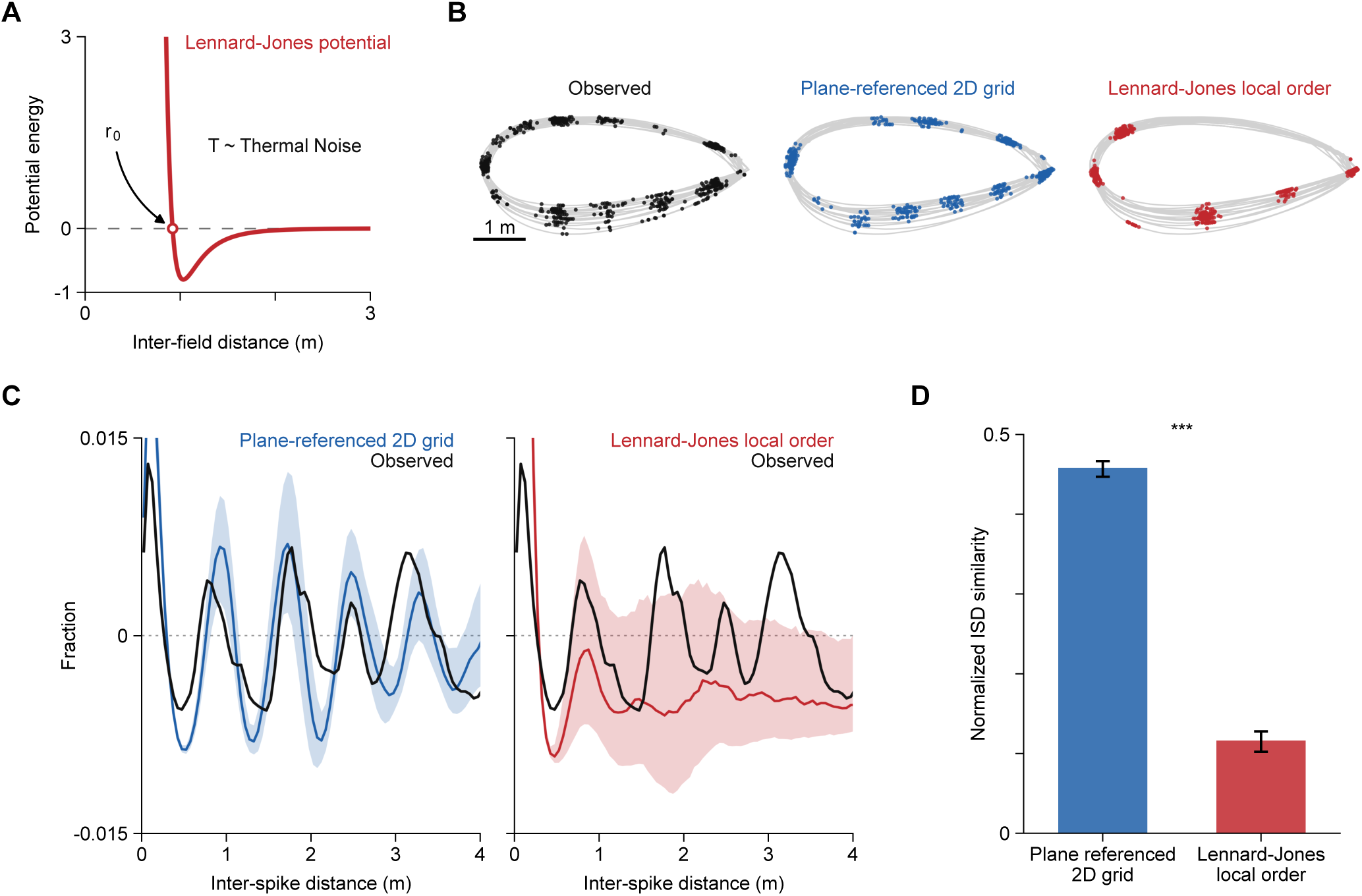
Lennard-Jones local order model predicts local structure but not global periodicity on clustered flights. A local order grid cell model, where inter-field-distances are governed by the Lennard-Jones (L-J) potential in volumetric space, has been previously proposed to explain the presence of local order but no global periodicity when all flights were pooled^18^. (**A**) Schematic of the L-J potential function. The L-J potential has two parameters: r_0_, which controls the characteristic distance between fields under equilibrium, and T, the temperature, which determines the magnitude of thermal noise. (**B**-**D**) We thus sought to determine whether the L-J local order model could also predict periodic firing patterns on clustered flights. We found that the local order model yielded field arrangements with consistent nearest neighbor distances, as expected from minimization of L-J potential energy around r_0_. However, the L-J local order model largely underpredicted the higher order periodic structure observed on clustered flights. (**B**) Spikes of an example cell plotted on a clustered flight path. Left, observed spikes. Middle, simulated spikes (blue) under the plane-referenced 2D grid model. Right, simulated spikes (red) under the L-J local order model. Simulated spikes are drawn from the 50^th^ percentile of best-fit field arrangements for each model. (**C**) Normalized inter-spike distance distributions (ISD) of an example cell. Shown ISD is baseline-subtracted by the ISD with uniform spike rate on the same occupancy for visualization. Simulated ISD under plane-referenced grid model (left, blue), and L-J local order model (right, red), are overlaid on the observed ISD (black). Shaded region indicates the inter-quartile range of sampled ISDs. Simulated ISD traces are the median ISD drawn from each fitted model. (**D**) Median normalized similarity between observed ISD and simulated ISD under the plane-referenced 2D grid model (left) and L-J local order model (right). Error bars indicate bootstrapped standard error. \*\*\**P* = 3.88 × 10^−38^, Wilcoxon signed-rank test, *W* = 1098, *n* = 255.

**Figure S17.**
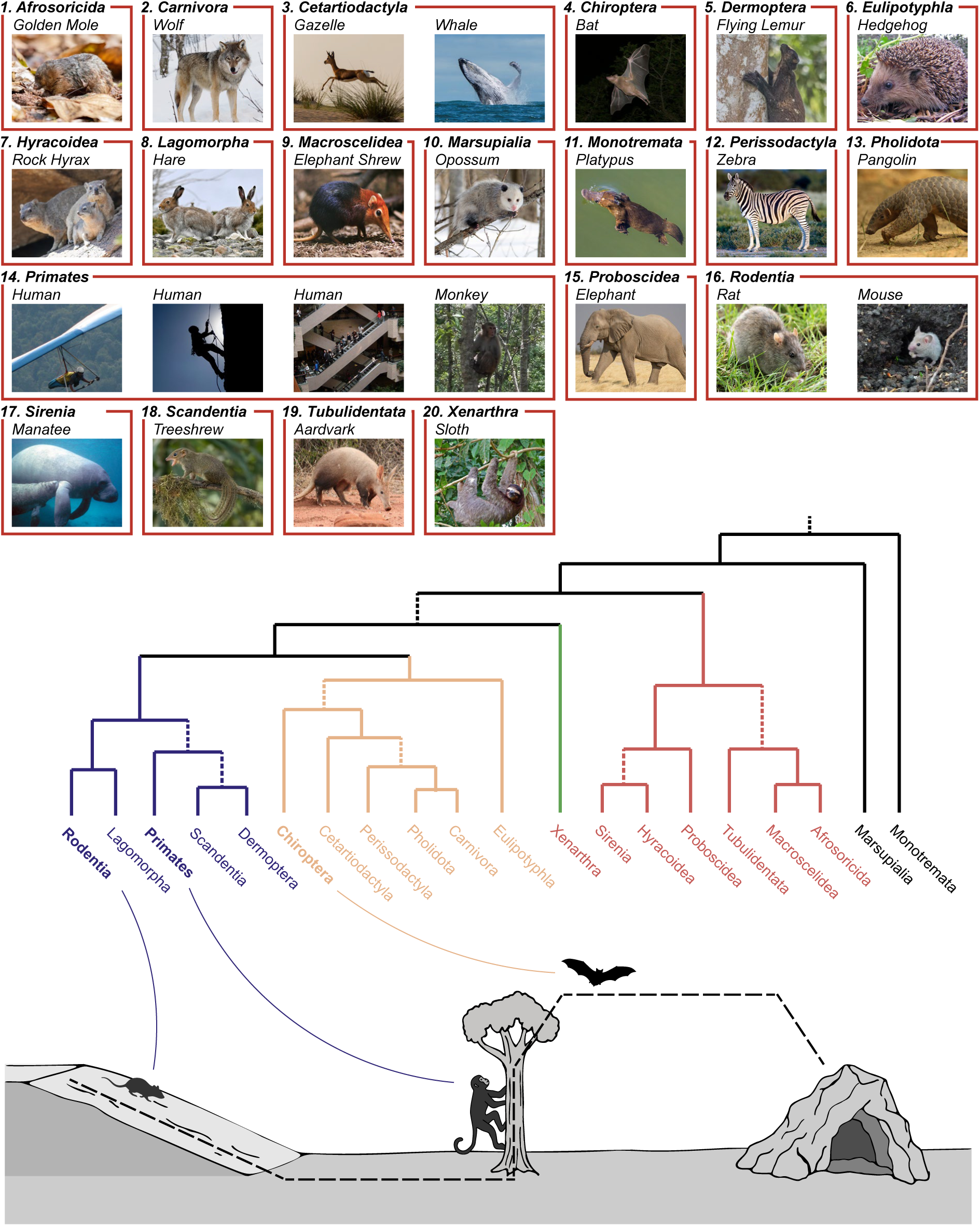
Behaviorally relevant movement subspaces in three-dimensional environments across mammals. Top, example species from 20 major mammalian lineages, illustrating the diversity of mammalian ecological niches and locomotor strategies. Middle, evolutionary tree of these lineages, modified and adapted from Springer et al.^86^. Bottom, schematic examples illustrating how movement through three-dimensional environments can often be organized along locally planar subspaces defined by species-specific ecology, body plan, and mode of locomotion, including terrestrial travel along ground surfaces, arboreal movement along trunks and branches, and aerial flight through structured planes of motion. These examples illustrate that, although mammals inhabit three-dimensional worlds, behavior and environmental geometry often structure movement along lower-dimensional movement manifolds.

**Table S1.** Persistent cohomology barcode statistics for all grid modules. Statistics of toroidal barcode features for *n* = 5 modules across flight and non-flight behavioral states in *n* = 3 bats. All modules exhibit significant toroidal signatures in at least one behavioral state (*n* = 3 bats; 4 of 5 with Bonferroni correction across states, *n* = 2 bats). Three out of five modules exhibit significant toroidal signature in both behavioral states (*n* = 2 bats). *n* = 500 shuffles. Torus detection is strongly dependent on recorded module size, and torus detection rates were similar to those reported in previous studies in rodents with similar module sizes^14^.

| Animal | Module | Behavioral State | H<sub>1</sub> (1D cycles) |  |  | H<sub>2</sub> (2D cavity) |  |  |
|---|---|---|---|---|---|---|---|---|
|  |  |  | Lifetime | P95 | P value | Lifetime | P95 | P value |
| Bat P | P1 | Flight | *3.36* | *1.93* | *P* < 0.002 | *3.95* | *2.68* | *P* < 0.002 |
|  |  | Non-flight | *3.62* | *3.38* | *P* = 0.018 | *5.25* | *4.66* | *P* = 0.012 |
|  | P2 | Flight | *3.17* | *2.57* | *P* = 0.006 | *3.54* | *3.47* | *P* = 0.040 |
|  |  | Non-flight | *4.79* | *3.78* | *P* = 0.016 | *6.47* | *5.82* | *P* = 0.022 |
|  | P3 | Flight | *3.11* | *2.33* | *P* = 0.006 | *5.14* | *3.01* | *P* < 0.002 |
|  |  | Non-flight | *3.62* | *3.41* | *P* = 0.020 | *4.55* | *4.94* | *P* = 0.100 (NS) |
| Bat Q | Q1 | Flight | *2.90* | *2.50* | *P* = 0.008 | *3.38* | *3.48* | *P* = 0.066 (NS) |
|  |  | Non-flight | *2.97* | *2.85* | *P* = 0.028 | *3.60* | *3.44* | *P* = 0.024 |
| Bat R | R1 | Flight | *2.94* | *2.12* | *P* < 0.002 | *3.48* | *3.14* | *P* = 0.010 |
|  |  | Non-flight | *3.95* | *3.64* | *P* = 0.008 | *4.11* | *3.92* | *P* = 0.026 |

